# Gene repertoire expansion and cis-regulatory diversification shaped 26S proteasome evolution within a proteostasis-centered network

**DOI:** 10.64898/2026.09.09.750301

**Authors:** Gautier Langin, Thomas Wilczek, David Biermann, Elizabeth A Rehmann, Evan S Forsythe, Suayb Üstün

**Affiliations:** Faculty of Biology & Biotechnology, Ruhr-University Bochum, Bochum, Germany; Institute of Molecular Plant Biology, BOKU University, Vienna, Austria; Center for Plant Molecular Biology (ZMBP), University of Tübingen, Tübingen, Germany; Biochemistry and Molecular Biology Program, Oregon State University-Cascades, Bend, OR, USA; Biology Program, Oregon State University-Cascades, Bend, OR, USA; Department of Integrative Biology, Oregon State University, Corvallis, OR, USA

**Author notes:** The author responsible for distribution of materials integral to the findings presented in this article in accordance with the policy described in the Instructions for Authors (https://academic.oup.com/plcell/pages/General-Instructions) is Gautier Langin.

## Abstract

The 26S proteasome is essential for proteostasis and constitutes one of the most conserved molecular machineries in eukaryotes. Its homeostasis is maintained by a mechanistically conserved feedback loop involving kingdom-specific components. Yet, how the proteasome subunits and associated regulatory feedback loop have evolved to accommodate ever-changing cellular context is poorly understood. Here, we combine gene duplicate analysis, cis-regulatory element identification, functional validation, and protein network inference to investigate the evolution of plant 26S proteasome. Proteasome gene repertoires expanded largely independently across plant lineages, with paralogs showing shifts in post-translational modification sites and pronounced transcriptional divergence in response to environmental cues. Comparative analysis of 26S proteasome gene promoters revealed extensive diversification of proteasome-associated cis-elements across plants, with repeated enrichment of related motifs. These observations led to the identification of telomere repeat-binding proteins (TRBs) as novel transcriptional regulators of proteasome genes in *A. thaliana*, through association with the previously characterized PRCE motif. Finally, evolutionary rate covariation analysis identified a conserved proteasome-associated network connected through proteasome-associated cis-elements and unifying cellular proteostasis. Together, our results indicate that plant 26S proteasome is controlled by a regulatory architecture that combines conserved cis-regulatory mechanisms, lineage-specific innovation, and stress-responsive paralog specialization at the center of the proteostasis network; offering a new perspective on the evolution of one of the most essential molecular complexes.

## Introduction

Most essential molecular processes that regulate eukaryotic life were already present in the last eukaryotic common ancestor (LECA) (Richards et al., 2024). Among these, pathways coordinating cellular proteostasis – from translation to protein degradation – are among the most conserved (Powers and Balch, 2013; Cox et al., 2026). Conversely, biological diversity among eukaryotes is shaped by the emergence of lineage-specific processes (Katz, 2012). Consequently, these highly conserved machineries had to accommodate novel molecular mechanisms while maintaining ancestral functions. The 26S proteasome complex exemplifies such a conserved system facing distinct constraints across eukaryotic lineages.

The 26S proteasome is a eukaryotic multi-heteromeric holoenzyme responsible for up to 80% of protein degradation in the cell (Collins and Goldberg, 2017). It evolved from prokaryotic proteasomes (Volker and Lupas, 2002). Its 20S core particle (CP), structurally related to the archaeal 20S proteasome, comprises two heptameric rings (alpha and beta) stacked to form a symmetric, four-layered proteolytic cylinder. The major difference from the archaeal proteasome is the presence of distinct subunits within each ring (Volker and Lupas, 2002). The 20S CP is capped by a heterohexameric AAA-type ATPase ring that controls gate opening. To form the 26S proteasome, this ATPase ring serves as a base to associate with additional regulatory particle (RP) subunits, providing a substrate-recognition platform (Wollenberg and Swaffield, 2001). Substrate recognition is mediated by ubiquitin receptors that recognize proteins modified by the ubiquitination machinery for selective degradation (Marshall and Vierstra, 2019). Together, the ubiquitination machinery and the 26S proteasome constitute the ubiquitin–proteasome system (UPS) (Marshall and Vierstra, 2019).

Although the UPS is highly conserved, notable organismal differences exist. Full assembly of the 26S proteasome in eukaryotes requires 31 distinct subunits, each of which is represented as a single-copy gene in the *Saccharomyces cerevisiae* genome (Hilt and Wolf, 1995). *Homo sapiens,* on the other hand, harbors specific duplications that support context-specific assembly (Murata et al., 2018) and *Arabidopsis thaliana* contains numerous duplicated subunits (Gladman et al., 2016). In *H. sapiens*, duplications act in a cell type- or stress-specific manner (Murata et al., 2018). While there is evidence that duplication of UPS subunits is linked to proteasome transcriptional regulation in *A. thaliana* (Gladman et al., 2016), the overall extent and functional implications of UPS gene duplication in plants are unknown, owing to the absence of up-to-date comparative genomic studies focusing on 26S proteasome evolution across plants.

Proteasome transcription is governed by a conserved regulatory feedback loop (Marshall and Vierstra, 2019; Langin et al., 2026). In all three major eukaryotic kingdoms, a proteasome transcriptional activator is itself a 26S proteasome substrate. Under proteasomal stress, degradation of the activator is attenuated, leading to its accumulation, nuclear translocation, and activation of proteasome gene expression (Krämer et al., 2021; Ruvkun and Lehrbach, 2023; Langin et al., 2026). Although this mode of regulatory feedback is conserved, neither the transcriptional activators nor the targeted cis-regulatory elements appear to share a common evolutionary origin. In *H. sapiens*, the bZIP transcription factors Nrf1/2 activate proteasome genes via the antioxidant response element (ARE) (Ruvkun and Lehrbach, 2023). In *S. cerevisiae*, the C2H2 transcription factor Rpn4 binds the proteasome-associated control element (PACE) (Mannhaupt et al., 1999). In *A. thaliana*, the plant-specific NAC53/78 factors activate proteasome genes via the proteasome-related cis-element (PRCE) (Nguyen et al., 2013; Langin et al., 2026).

In *A. thaliana*, response to proteasome perturbation by chemicals and pathogens shows a bias among proteasome paralogs transcriptional activation (Gladman et al., 2016; Langin et al., 2026). Proteasome transcriptional regulation is also linked to other essential cellular processes, including energy metabolism (Krämer et al., 2021; Langin et al., 2026). These observations suggest that proteasome gene duplication contributes to maintenance of plant 26S proteasome homeostasis and that the feedback loop connects the proteasome to broader gene regulatory networks. However, the origins of these duplications, the extent of divergence between paralogs, and the conservation of the plant proteasome transcriptional regulatory module remain unclear. Detailed analysis are needed to clarify proteasome regulation in plants, to define its integration into proteostasis regulatory networks and the larger cellular interactome, and to determine how this highly conserved molecular complex adapts to an ever-changing cellular proteome.

Here, we performed a comparative genomic analysis of 18 representative Viridiplantae species spanning from chlorophyte algae to angiosperms. We analyzed the origins of proteasome gene duplications, their evolution across plants, and divergence between paralogs. We found that each species harbors a specific set of proteasome gene duplicates, where diversification of proteasome subunits is associated with fine-tuning of electrostatic potential and divergence in post-translational modification sites. We further show that proteasome paralogs are differentially transcriptionally regulated by environmental cues in both angiosperms and distantly-related Viridiplantae lineages. Moreover, promoter analysis identified novel proteasome-associated cis-elements across plants, including significant enrichment of six distinct motifs that differ across the 18 study species, highlighting striking diversification of proteasome-associated cis-regulatory architecture. We validated conservation of the NAC53/78 regulatory module in angiosperms and found that telomere repeat–binding proteins (TRB) can recognize proteasome-associated cis-elements from eight species, from the streptophyte algae *Chlorokybus atmophyticus* to mesangiosperms. Consistent with this, telomere repeat–binding proteins contribute to proteasome homeostasis maintenance in *A. thaliana*. Finally, we show that the plant 26S proteasome is embedded in a protein network centered on proteostasis regulation and connected by proteasome-associated cis-elements specific enrichment throughout plant evolution. Overall, our findings provide a new view of 26S proteasome evolution in plants, uncover ancient regulatory factors, and highlight a conserved proteostasis-centered network linked by proteasome-associated cis-regulatory control.

## Materials and methods

### Genomic Data

We used the following genome releases: *Saccharomyces cerevisiae* Ensembl R64-1-1 (SACC); *Chlamydomonas reinhardtii* JGI CC-503 v5.6 (CHRE); *Chlorokybus atmophyticus* JGI CCAC 0220 (CHAT); *Zygnema circumcarinatum* JGI SAG 698-1b (ZYCI); *Zygnema cylindricum* JGI SAG 698-1a (ZYCY); *Anthoceros agrestis* AagrOXF (ANAG); *Marchantia polymorpha* Tak v6.1r2 (MAPO); *Physcomitrium patens* JGI v6.1 (PPAT); *Selaginella moellendorfii* NCBIrefSeq (SELM); *Diphasiastrum complanatum* JGI v3.1 (DICO); *Ceratopteris richardii* JGI v2.1 (CERI); *Gingko biloba* v1 (GBIL); *Cryptomeria japonica* SUGI v1 (CRJA); *Amborella trichopoda* JGI 75 HAP1 v2.1 (AMTR); *Oryza sativa* Ensembl IRGSP-1.0 (OSAT); *Solanum lycopersicum* ITAG4.1 (SOLY); *Arabidopsis thaliana* TAIR10 (ARAT); *Populus trichocarpa* Ensembl v4.1 (POTR); *Glycine max* Ensembl Wm82 a4_v2.1 (GLYM).

### Promoter sequences

Promoter sequences were extracted from GFF annotations using the promoterExtract Python package (v0.9.5.8), with promoter length set to 2000 bp or 400 bp upstream of the TSS and the 5′ UTR extension set to 100 bp downstream of the TSS (i.e., −2000/+100 or −400/+100 windows relative to the TSS).

### Protein sequences

For protein-level analyses, we used the representative/primary transcript per gene as designated in each genome release. If no representative transcript was annotated, one transcript per gene was retained for downstream analyses.

### Orthologs identification

To identify genes encoding 26S proteasome subunits, *Saccharomyces cerevisiae* proteasome subunit sequences were queried against plant proteomes using HMMER v3.3.2. Candidate hits were filtered and manually curated to remove false positives (**Table S1**).

### Cis-regulatory elements analysis

Cis-elements associated with proteasome promoters were identified by pattern matching implemented in R. We generated, *in silico*, a comprehensive motif library comprising: (i) all IUPAC-encoded nucleotide motifs of length 4–7 bp; (ii) motifs of length 8–9 bp allowing one ambiguous position; and (iii) gapped motifs with internal spacer lengths of 7–12 bp flanked by 3–5 bp on each side. These patterns were matched to promoter sequences using regular-expression functions.

Motif counts were obtained for three promoter windows: −400/+100 bp relative to the TSS for all genes, −400/+100 bp for 26S proteasome genes, and −2000/−400 bp for 26S proteasome genes. Genome-wide enrichment was assessed by Fisher’s exact test comparing motif counts in proteasome promoters (−400/+100) *versus* all promoters (−400/+100). Positional bias was assessed by Fisher’s exact test comparing motif counts in proteasome proximal promoters (−400/+100) versus distal promoter regions (−2000/−400) of the same genes. Motifs that were under-represented (present in <50% of proteasome promoters within the −400/+100 window) or strongly over-represented (>300% enrichment in proteasome promoters within −400/+100) were filtered out. For each motif, we summed the negative log10–transformed false discovery rates from the genome-wide enrichment and positional-bias tests to obtain a composite score.

These patterns were then matched against promoter sequences. Counts were generated against promoter sequences ranging from -400bp from TSS to +100bp of all genes, against - 400bp/+100bp of 26S proteasome genes promoters and -2000bp/-400bp of 26S proteasome genes promoters.

Genome wide enrichment was assessed through fisher exact test of 26S proteasome promoters -400bp/+100bp counts versus all promoters -400bp/+100bp counts. Positional biases were assessed through fisher exact test of 26S proteasome promoters -400bp/+100bp counts versus 26S proteasome promoters -2000bp/-400bp counts. Unrepresented motifs (less than 50% frequency in 26S promoters -400bp/+100bp) and over-represented motifs (more than 300% enrichment in 26S promoters -400bp/+100bp) were filtered out. Negative log10 transformation of false-discovery rates for fisher exact tests for genome wide enrichment and positional biases were added together to compute a score for each motif.

For positional visualization, the selected motifs were scanned by regular expression across −2000/+100 bp promoter sequences. Motif frequency was computed in 20 bp bins and displayed as bar plots; kernel density curves were overlaid where helpful to illustrate positional enrichment.

For motif enrichment analyses in promoters (−400/+100 bp) of ERC co-evolution network members, we used STREME (MEME suite v5.5.9) (Bailey et al., 2015) with default parameters, providing all gene promoters (−400/+100 bp) from the corresponding genome as the background/control set.

### Multiple-sequence alignment

For analyses of 26S proteasome subunit evolution and sequence constraints, codon-based multiple sequence alignments were generated with MACSE v2.0.7 (Ranwez et al., 2021). Each subunit family was aligned independently using the corresponding coding sequences (CDS).

Protein sequences of NAC53/78-related transcription factors were aligned with Clustal Omega (version as installed at run time). The sequence set included all *A. thaliana* NAC transcription factors bearing an N-terminal transmembrane (NTM/NTL) motif, the reference NAC AtANAC019, the closest NAC78 ortholog from *M. polymorpha* (MpNAC4), and top angiosperm matches identified by BLASTp searches using the *A. thaliana* NAC78 protein as the query. BLASTp searches were run against angiosperm proteomes with default parameters; top hits were retained per species.

### Phylogenetic analysis

For phylogenetic inference of 26S proteasome subunits, nucleotide alignments generated with MACSE v2.0.7 were analyzed with IQtree v2.3.6 (Minh et al., 2020). Support was assessed using 1,000 SH-like approximate likelihood ratio test (SH-aLRT) replicates and 1,000 bootstrap replicates. The resulting consensus gene trees were reconciled with the species tree using Notung v3.0 (Chen et al., 2000). The species tree was derived from NCBI taxonomy (Schoch et al., 2020). Proteasome subunit trees were rooted and rearranged under Notung’s default parameters, including a support threshold set to 90% of the highest edge weight in the gene tree. To reconstruct the history of proteasome subunit duplications in plants, rearranged gene trees were inspected and inferred duplication and loss events were mapped onto the species tree.

Positive selection analyses used the MACSE nucleotide alignments and PAML4.9j (with PAMLX v1.3.1) (Álvarez-Carretero et al., 2023). We applied the branch (free-ratio) model with one ω estimated per branch. Branches were excluded from downstream interpretation if they showed unstable estimates dN/dS > 10 or effectively zero divergence (dN < 1e−5) coupled with dN/dS > 1. For comparisons between paralogs, we considered ω estimates on branches immediately following duplication events.

For NAC53/78-related transcription factors, a protein tree was generated from the Clustal Omega alignment, using the program’s HMM profile–based guide tree as the topology.

### Protein sequence analysis: IDR, PTM sites, Entropy & Electrostatic charge

For protein sequence analysis of 26S proteasome subunits, amino acids sequence alignments from MACSE were for residue position determination and other analysis.

Intrinsically disordered region score for representative sequences was calculated using AIupred algorithm with default parameters (Erdős and Dosztányi, 2024).

Local electrostatic charge was calculated using idpr R package with an amino acid window of 9 residues (McFadden and Yanowitz, 2022).

Hierarchical clustering was performed using average prediction from subunit paralogs for every species. Based on multi-sequence alignment, only predictions at positions with less than 25% missing residues were considered. For every subunit family, a distance matrix was built based on predictions at retained positions. For every subunit class, an average distance matrix was built from individual distance matrix. Based on the average distance matrix, hierarchical clustering was performed using the method “average” in R and a dendrogram was used to represent the results.

Local Electrostatic Charge Variance between paralogs was calculated as the average of the local electrostatic charge variance at every position between paralogs and visualized with a heatmap.

Shannon entropy for residue position in proteasome subunits alignments was calculated using entropy function from Bios2cor R package with a gap ratio of 80%.

For post-translational modification site analysis, sites were extracted from the database qPTMplants (Xue et al., 2022) for the species *Chlamydomonas reinhardtii*, *Physcomitrium patens*, *Glycine max*, *Solanum lycopersicum*, *Populus trichocarpa*, *Arabidopsis thaliana* and *Oryza sativa subsp. Japonica.* For each PTM site, variability at the corresponding alignment position was analyzed. Percentage of residue conservation was caculated as the proportion of sequences possessing the same residue as the one identified to be modified. Because phosphorylation can happen on several residues, for this PTM, substitution between S, T and Y was considered as conservation.

### *In silico* 20S CP reconstitution

To reconstitute 20S CP *in silico*, AlphaFold models for every 26S proteasome subunits from *C. reinhardtii*, *M. polymorpha* and *A. thaliana* were downloaded. Using ChimeraX 1.11 (Pettersen et al., 2021), the crystal structure of *S. cerevisiae* (PDB: 1RYP) was used as template and every plant species Alphafold subunit models (Jumper et al., 2021) wassuperimposed to the corresponding *S. cerevisiae* subunit. After complete reconstitution, coloring based on the Coulombic electrostatic potential was used to color molecular surfaces.

### Plant material, growth and treatment

Arabidopsis thaliana plants were grown on ½ Murashige and Skoog (MS) solid medium with 0.8% agar plus 1% sucrose. Seeds were surface sterilized 10min with 1.3% sodium hypochlorite and stratified for 2 days after sawing. Plants were grown under long day conditions (light/dark cycles: 16h 22°C/8h 20°C, 130µmol.mm².s^-1^ light intensity, 60% relative humidity). For phenotyping, solid medium was supplemented with indicated amount of bortezomib or mock solution. After 10 to 14 days of growth, fresh weight was measured. For one replicate, 5 representative seedlings were scaled together on an analytical scale (resolution 0.0001g). For mRNA expression or western blot analysis, 7 days after germination seedlings were transferred to liquid ½ Murashige and Skoog (MS) plus 1% sucrose supplemented with indicated concentrations of bortezomib or mock solution for 6 hours prior sampling.

*Marchantia polymorpha* plants were obtained from the Ebert Lab and grown on a ½ Gamborg B5 solid media with 1% agar in a 16 h light and 8 h dark cycle with a light intensity of 120 mmol m^-2^ s^-1^ at 21 °C and constant relative humidity of 70%. For western blot analysis, *Marchantia* plants were transferred to liquid ½ Murashige and Skoog (MS) plus 1% sucrose supplemented with indicated concentrations of bortezomib or mock and subjected to vacuum infiltration 2 times 1 min at +/- 13 mBar and incubated for 6 hours prior sampling.

*Selaginella kraussiana* plants were obtained from Ruhr-University Bochum botanic garden. For western blot analysis, *Selaginella* tissues were transferred to liquid ½ Murashige and Skoog (MS) plus 1% sucrose supplemented with indicated concentrations of bortezomib or mock and subjected to vacuum infiltration 2 times 1 min at +/- 13 mBar and incubated for 6 hours prior sampling.

### RNA isolation & RT-qPCR analysis

Total RNA isolation was performed using NulceoZOL (MACHEREY-NAGEL 740404.200) according to manufacturer instructions. To exclude potential contaminant DNA, RNA samples were subjected to DNase I treatment (Thermo scientific™) following provider instructions. For RNA sequencing, integrity and RNA concentration was determined (2100 Bioanalyzer, Agilent). For RT-qPCR sample analysis was performed as described previously; cDNA synthesis was performed using LunaScript® RT SuperMix Kit (NEB) following provider recommendation. qPCR was performed using MESA BLUE qPCR 2X MasterMix Plus for SYBR® Assay (Eurogentec) using a 2-step reaction protocol for 40cycles with systematic evaluation of primer melting curve. mRNA level was quantified based on the ΔΔCt method followed by Log2 transformation.

### Immunoblot analysis

For immunoblot analysis, sample processing was performed as described previously. Plant tissue was homogenized in protein extraction buffer (100mM Tris-HCl pH 7.5, 3% SDS, 10mM EDTA). Protein extracts were boiled with Laemmli buffer for 10min at 95°C and centrifuged for 1min at 13000g, supernatants were used for SDS-PAGE.

Western blotting was done by SDS-PAGE using TGX FastCast 10% Acrylamide gels (Biorad). SDS-PAGE was performed in TGS running buffer (25 mM Tris pH 8.3, 192mM glycine, 0.1% SDS) and proteins were transferred on PVDF membrane 0.2mm. PVDF membranes were incubated for 1h in TBST (50mM Tris-HCl pH 7.5, 150mM NaCl, 0.1% Tween20) solution containing 5% Skimmed-Milk followed by antibody incubation in TBST 5% Milk for 1h at RT or over-night at 4°C. Membranes were washed 3 times in TBST for 5min and 1 time in TBS for 5min. HRP chemiluminescence was detected using Amersham™ ECL Select™ detection reagent and images were taken using a CCD camera.

For signal quantification, raw images of the immunoblot signals against the protein of interest and the corresponding loading control were processed using the dedicated software Image Lab Software (https://www.bio-rad.com/de-at/product/image-lab-software?ID=KRE6P5E8Z).

### Transcriptomic analysis

For global and tissue specific expression level analysis, FPKM for *A. thaliana*, *G. max* and *O. sativa* were obtained from https://plantrnadb.com (Yu et al., 2022), using only libraries annotated as untreated. RPKM for *P. patens* were obtained from gene atlas previously published (Perroud et al., 2018). TPM for *S. moellendorfii* were calculated from previously published expression atlas (Ferrari et al., 2020). For correlation analysis between paralogs across tissues/developmental stages, Spearman correlation index was calculated using average counts values for every paralog in given tissue/stage.

For transcriptome analysis under environmental perturbation, raw FastQ files from several datasets were downloaded. FastQ files were subjected to preprocessing and quality control with fastp python package (Chen, 2025) following developer recommendation. TPM and estimated counts were computed with kallisto software (Bray et al., 2016) following developer recommendations. Used datasets are detailed in **Table S2**.

For differential expression analysis, estimated counts were used with the R package DEseq2 (Love et al., 2014). Genes were considered as differentially expressed if false discovery rate < 0.05 and Log2 fold change were used for heatmap representation.

For comparison between *A. thaliana* subunits expression in response to environmental perturbation, Spearman correlation of expression fold-change was calculated between non-duplicated subunits genes or between paralogs pairs.

For paralog bias analysis, transcriptome data were categorized by context and for every species the absolution maximum difference between Log2 fold change values between paralogs was calculated.

For co-expression analysis of ERC network members with proteasome genes, TPM calculated from the datasets analyzed in this study (**Table S2**) for the respective species were used. Median of the spearman correlation indexes was calculated for every network member against all proteasome genes using only correlations with p value < 0.05. For negative control, 3 independent sets of genes of the same size as the network genes were used. For positive control 26S proteasome genes were used.

For analysis of paralogs differences in co-expression with ERC network members, absolute differences between Spearman correlation index for proteasome gene paralogs against all ERC network members were calculated. For control, median differences of Spearman correlation index for all non-duplicated proteasome genes against all ERC network members were calculated.

### Protein purification

Recombinant proteins were expressed in E. coli BL21(DE3) by IPTG induction. Bacterial solution was centrifuged 20min at max speed and pellet was subjected to sonication after resuspension in 1mL MBP-buffer (20 mM TRIS-HCl pH 7.5, 200mM NaCl, 1mM EDTA). Sonicated samples were incubated with amylose resin (New England Biolabs) for MBP-tagged transcription factors binding. After 2h incubation on rotating wheel, tubes were centrifuged and supernatant removed. Proteins were eluted by resuspending the amylose resin in MBP-buffer with 50mM maltose and incubating for 30min.

### Electromobility Shift Assay

For EMSA assays, complementary DNA oligos were synthesized by EuroFins Genomics with ATTO565 dye linked to the 5’ end of minus strand for probes. To generate double stranded probes, complementary oligos were incubated at 2µM in annealing buffer (25mM HEPES-KOH pH 7.8, 40mM KCl) at 70°C for 5min and cool down to room temperature.

EMSA reactions were performed in 20µL reaction buffer (25 mM HEPES-KOH pH 7.8, 40 mM KCl, 1 mM DTT, 10% Glycerol) using 1µg of purified MBP-NAC53/78 and 50ng DNA probe for 30min at 25°C. Reactions were subjected to native polyacrylamide migration with TGX FastCast 7.5% Acrylamide gels (Biorad) in TAE (Tris-base 40 mM pH 8.3, 20mM acetic acid, 1mM EDTA). After migration, probes were imaged using a BioRad Chemidoc ™ Imaging System. For competitor assay, unlabeled mutated probed were added 10min before the end of the reaction at the indicated concentration.

For shift quantification, BioRad Image Lab Software was used to quantify the ratio between the signals of the shifted and unshifted bands.

### ChIP-Seq and ATAC-seq data analysis

To analyze *in vivo* association of TRBs with proteasome promoters, we mapped ChIP-seq signal of tagged TRB1, TRB2, TRB3 and TRB4 protein from *Arabidopsis thaliana* (Wang et al., 2023; Amiard et al., 2024) (**Table S2**). We mapped average signal from experimental replicates along the -3kb/+3kb around proteasome genes TSS compared to signals in respective control conditions.

To analyze chromatin accessibility at proteasome promoters, we examined significantly open chromatin region in the 26S proteasome promoter regions -2kb/+100bp from previously published ATAC-seq on *A. thaliana* upon bacterial infection (Ding et al., 2021).

To correlate TRB ChIP-seq signal with PRCE presence or chromatin accessibility, we analyze signal in region containing the GCCCA core motif or within significantly open chromatin regions in the 26S proteasome promoter regions and compared to control conditions signal.

### ERCnet

We input our proteome fasta files into Orthofinder v.2.5.5 (Emms and Kelly, 2019) to cluster homologous gene families and applied ERCnet v1.3.0 (Forsythe et al., 2025) to the gene families. ERCnet was run with default parameter values, except the values specified below (see ERCnet documentation (https://github.com/EvanForsythe/ERCnet) for detailed parameter descriptions). The *Phylogenomics.py* step was run with -p=4, -r=10, and the -T option. The ERC_analyses.py step was run with -b=BXB. Finally, the Network_analyses.py step was run with -pp=0.01, -sp=0.01, kp=0.01, -pr=0.4, -sr=0.4, and -kr=0.3. These filtering threshold values equate to retaining pairs of genes as ERC hits if they meet the following requirements: Pearson’s P-value<0.01, Spearman’s P-value<0.01, Kendall’s P-value<0.01, Pearson’s R>0.4, Spearman’s R>0.4, and Kendall’s Tau>0.3. These thresholds were selected empirically by comparing the distribution of ERC scores for subunits of the Caseinolytic Protease complex, which were used as positive control based on prior results (Gatts et al., 2026), to the genome-wide background distribution (Fig S16). The final thresholds were chosen to provide good separation between these distributions.

Proteins with significant ERC hits according to our thresholds constituted the nodes of our predicted interaction network. We visualized our network and performed downstream analyses with Cytoscape (Shannon et al.). We integrated *A.thaliana* interaction predictions from the STRING database (Szklarczyk et al., 2023) by color coding existing ERC-based edges that also have STRING-based score greater than 150. Additionally, we added new STRING-based edges among pairs of 26S proteasome nodes. Candidate interaction nodes were colored according to select enriched GO categories. We focused on the subnetwork containing 26S proteasome nodes by extracting the nearest neighbor nodes to all 26S proteasome nodes, yielding our 26S proteasome interaction network (Fig 5A). These direct ERC hits with 26S proteasome units were used for downstream functional enrichment analyses. We applied the Glay clustering algorithm to visualize clustering structure within the subnetwork. For downstream analysis, all paralogs for all nodes were considered in the respective plant species.

### Gene ontology

For Gene Ontology enrichment, gene IDs from A. thaliana genome corresponding to the ERC network members were provided to ShinyGOv0.85 (Ge et al., 2020). Top50 enriched terms for Cellular Components and Biological Process were considered with FDR cutoff of 0.05 and visualized by dendrograms representation.

### Statistical Analysis

Statistical tests used to assess significance are detailed in the figure legends.

## Results

### 26S Proteasome gene repertoire expanded largely during green lineage evolution

To capture broad patterns of 26S proteasome regulatory features diverging across model eukaryotes, we compiled gene copy number counts and position in promoters of core proteasome-associated cis-elements from *S. cerevisiae*, *H. sapiens*, and *A. thaliana* (**Fig S1A, B**). Consistent with previous studies, we found that proteasome gene repertoires differ markedly across eukaryotes (**Fig S1A**) and known cis-elements from *S. cerevisiae* and *A. thaliana* are characterized by a strong positional bias in proximity to 26S proteasome genes transcription starting site (TSS) (**Fig S1B**), highlighting the importance of these two features in 26S proteasome evolution.

To assess the frequency of proteasome gene duplication across eukaryotic evolution, we queried the OMA orthology database for the number of duplicated subunits in >900 eukaryotic species (Altenhoff et al., 2024) and found that plant species showed the largest proteasome gene expansions (**Fig S2A**). Because gene family expansion often tracks genome expansion, especially in plants (Almeida-Silva and Van de Peer, 2023), we compared the number of duplicated subunits to total gene number. We observed a strong correlation between proteasome duplications and total gene number (**Fig S2B**), but this relationship was largely driven by species with very large gene counts. Restricting analysis to species with 5,000–35,000 genes substantially reduced the correlation (**Fig S2B**). Assessing this relationship by kingdoms further revealed substantial differences in the number of proteasome duplicates within comparable genome-size ranges in plants and animals (**Fig S2C**). These observations indicate that proteasome gene expansion is a recurrent feature in both plants and animals and is only partially explained by genome expansion.

To investigate proteasome evolution within the green lineage, we identified proteasome genes in 18 representative species based on sequence homology to *S. cerevisiae* subunits, spanning from chlorophytic algae to angiosperms. We found that the number of genes encoding 26S proteasome subunits varied considerably across species, and within monophyletic groups (**Fig 1A, Table S1**), suggesting multiple, lineage-specific expansion events. Examination of the repertoire within each species (**Fig S3**) identified five species with missing subunits, likely reflecting incomplete data in early-stage genome assemblies. We detected duplicate gene copies in 16/18 species, beginning with *C. reinhardtii* (**Fig S3**), indicating that duplications are not restricted to large genomes. Duplication patterns were species-specific, with closely related species often showing distinct profiles (**Fig S3**). Notably, alpha CP subunits, RPTs, and RPN11 were frequently duplicated, whereas RPN12 and Beta7 were comparatively resistant to duplication (**Fig S3**). These patterns suggest that most duplications do not share common origins and that individual subunits are subject to distinct evolutionary constraints.

**Figure 1.**
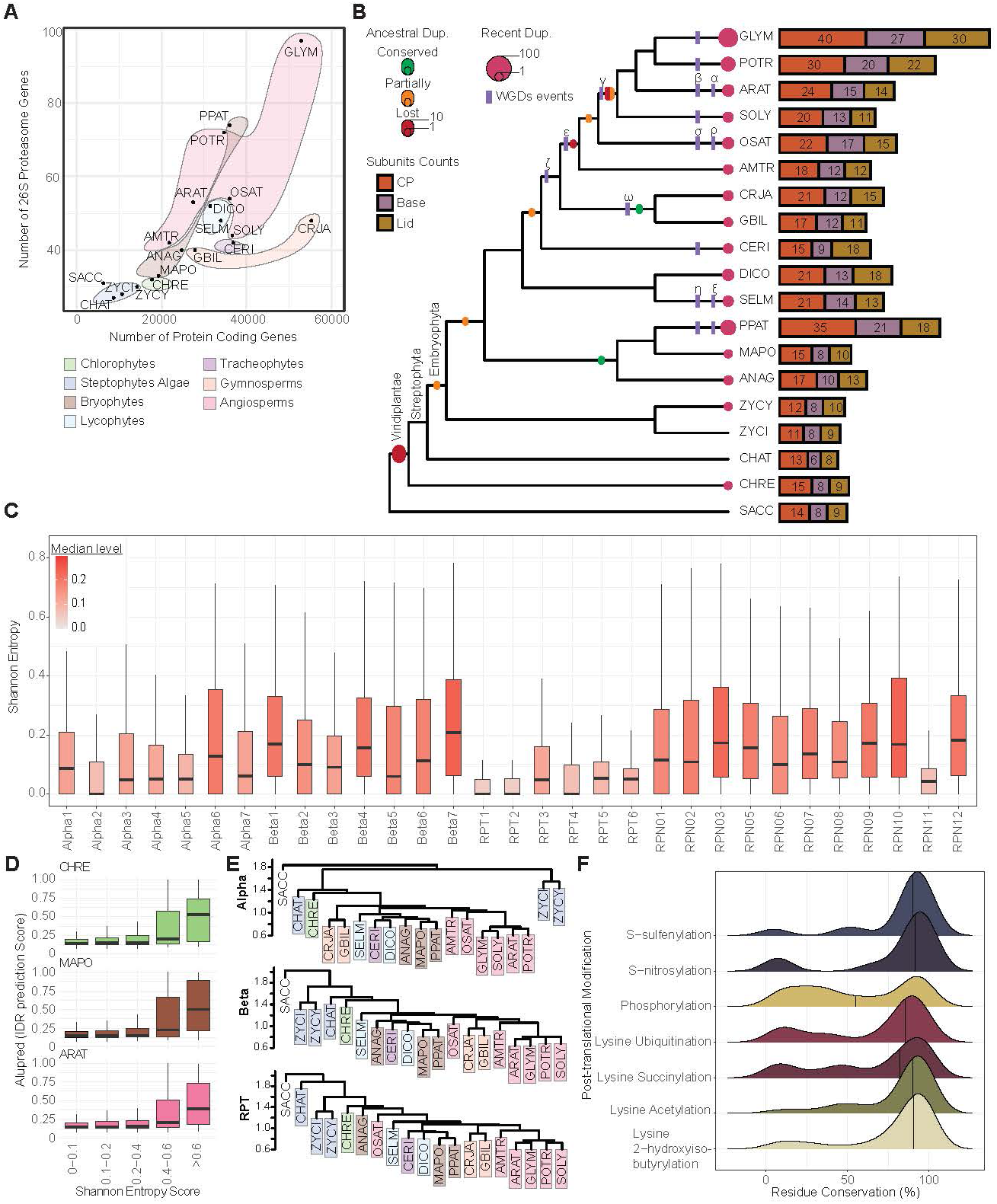
Plant 26S Proteasome diversification arose from gene repertoire expansion. **A.** Number of 26S proteasome genes versus total protein-coding genes across representative plant species; phyletic groups are indicated by colored areas. **B.** Origins of proteasome gene duplications mapped onto the plant species tree. Predicted proteasome gene duplications and documented whole-genome duplication (WGD) events are indicated. Stacked bar plots show the number of proteasome genes per subunit class for each species. **C.** Shannon entropy from multiple-sequence alignments (FigS4) for each proteasome subunit family. Boxplot color encodes the median entropy per family. **D.** Intrinsically disordered region (IDR) prediction scores for residues of proteasome subunits in C. reinhardtii, M. polymorpha, and A. thaliana, plotted by Shannon Entropy score at the corresponding alignment positions. **E.** Hierarchical clustering dendrograms based on predicted electrostatic potential for α CP subunits, β CP subunits, and RP base subunits. Colored boxes indicate phyletic groups as in A. The dendrograms Y-axis indicates clustering distance. **F.** Density plot of residue conservation across plants for the different post-translational modification types.

To infer the timing of these duplication events, we performed codon-based alignments followed by maximum-likelihood phylogenetic analyses and gene tree/species tree reconciliation for each subunit family (**Fig 1B**, **Fig S4**) (Chen et al., 2000; Minh et al., 2020; Ranwez et al., 2021). Most duplication events were recent, lineage-specific duplications (**Fig 1B**). Across the 31 proteasome subunits, we inferred 25 ancestral duplication events. Of these, only four yielded conserved duplicated genes, six were partially conserved, and 15 were followed by loss of one duplicate in all descendant species (**Fig 1B**). Mapping documented whole-genome duplications onto the trees indicated that ancestral WGDs had limited impact on proteasome gene expansion (**Fig 1B**), again indicating that expansion of proteasome gene families is not directly tied to expansion of the genome at large. Together, these data indicate that most proteasome duplicates emerged recently, producing lineage-specific subsets of paralogs and pointing to recurrent, specific tuning of the proteasome gene repertoire during plant evolution.

### Plant 26S proteasome subunits evolution is characterized by purifying selection and sequence fine tuning

Our analyses suggest that each species harbors a specific proteasome gene repertoire. To investigate this specificity, we examined proteasome subunit protein sequence evolution. We first quantified sequence diversification for each subunit gene family using Shannon entropy and found that entropy varied markedly among gene families (**Fig 1C**, **Fig S5**). RPT subunits forming the AAA-ATPase ring showed low variability among homologs. Likewise, α CP subunits were generally among the least variable, with the exception of α6. By contrast, β CP subunits were relatively variable despite their central role in proteolysis. Most RP subunits were highly variable, with the notable exception of RPN11. These patterns indicate that proteasome subunit families experienced different levels of sequence diversification during plant evolution.

To understand the basis of this variability, we combined entropy profiles with predictions of intrinsically disordered regions (IDRs) and electrostatic potential (**Fig S5**). The regions of highest variability were frequently confined to N- or C-terminal segments, correlating with predicted IDRs (**Fig 1D**, **Fig S5**), consistent with known evolutionary lability of IDRs (Singleton and Eisen, 2024). In addition to previously characterized IDRs, our analyses predicted novel IDRs such as in CP Alpha4, Alpha6, RPT2, RPT4, RPN1 and RPN8 subunits (**Fig S5**). We also observed substantial variability at positions predicted to be ordered (**Fig 1D**, **Fig S5**). We hypothesized that such changes might alter electrostatic potential. Hierarchical clustering based on predicted residue charge for α CP, β CP, and RPT families (**Fig 1E**, **Fig S5**) produced an evolutionary projection largely concordant with species relationships, with most species clustering with close relatives (**Fig 1E**), suggesting 26S proteasome subunits electrostatic potential has diverged through fine-tuning in plants. To visualize surface electrostatics and IDRs in context, we reconstituted the 20S proteasome *in silico* using AlphaFold models (Jumper et al., 2021). Surface potential differed markedly from yeast to plants (**Fig S6A**) and our novel α CP subunit IDRs appeared compatible with complex assembly (**Fig S6A**), making these IDRs a credible feature of the 20S proteosome structure. Within plants, we detected differences in electrostatic potential across disordered tails and exposed CP surfaces (**Fig S6A, B**), highlighting fine-scale surface adaptation.

Because post-translational modifications (PTMs) can modulate surface properties, complex assembly and activity (Hirano et al., 2016; Kors et al., 2019), we next analyzed the variability of reported PTM sites in plants (Xue et al., 2022). From 527 modified positions identified in at least one of seven species (**Table S3**), we assessed conservation across homologs (**Fig 1F**, **Fig S7A**). We found numerous PTM sites – particularly phosphorylation sites – were poorly conserved across homologs (**Fig 1F**, **Fig S7A**). Given that IDRs are often subjected to PTMs (Bah and Forman-Kay, 2016), we examined PTM conservation relative to predicted disorder (**Fig S5**), but found only a weak relationship, with only a small number of PTMs mapping to strongly disordered regions (**Fig S7B**). Thus, most PTMs occur in ordered regions, likely structurally conserved, among homologs.

These results support significant evolutionary change within a structurally conserved proteasome. We then asked whether gene family expansion drives elevated rates of protein evolution. Across families, higher duplication counts were associated with lower average entropy among homologs (**Fig 2A**), indicating that duplication has not driven sequence variability. To test divergence after duplication, we analyzed predicted electrostatic potential and PTM sites variability between paralogs (**Fig 2B, C, Fig S8**, **Table S3**). Consistent with our findings that both features have diverged across plant species, paralogs also displayed differences with frequent mutations of PTM sites and variability in predicted electrostatic potential between paralogs. To test for adaptative evolution between paralogs, we performed positive selection analyses focused on post-duplication branches (Álvarez-Carretero et al., 2023). Despite numerous duplications, we detected signals of positive selection in only four cases (**Fig S9**), whereas the remaining 882 genes showed very low dN/dS ratios (**Fig 2D**, **Fig S9**), indicative of strong purifying selection.

**Figure 2.**
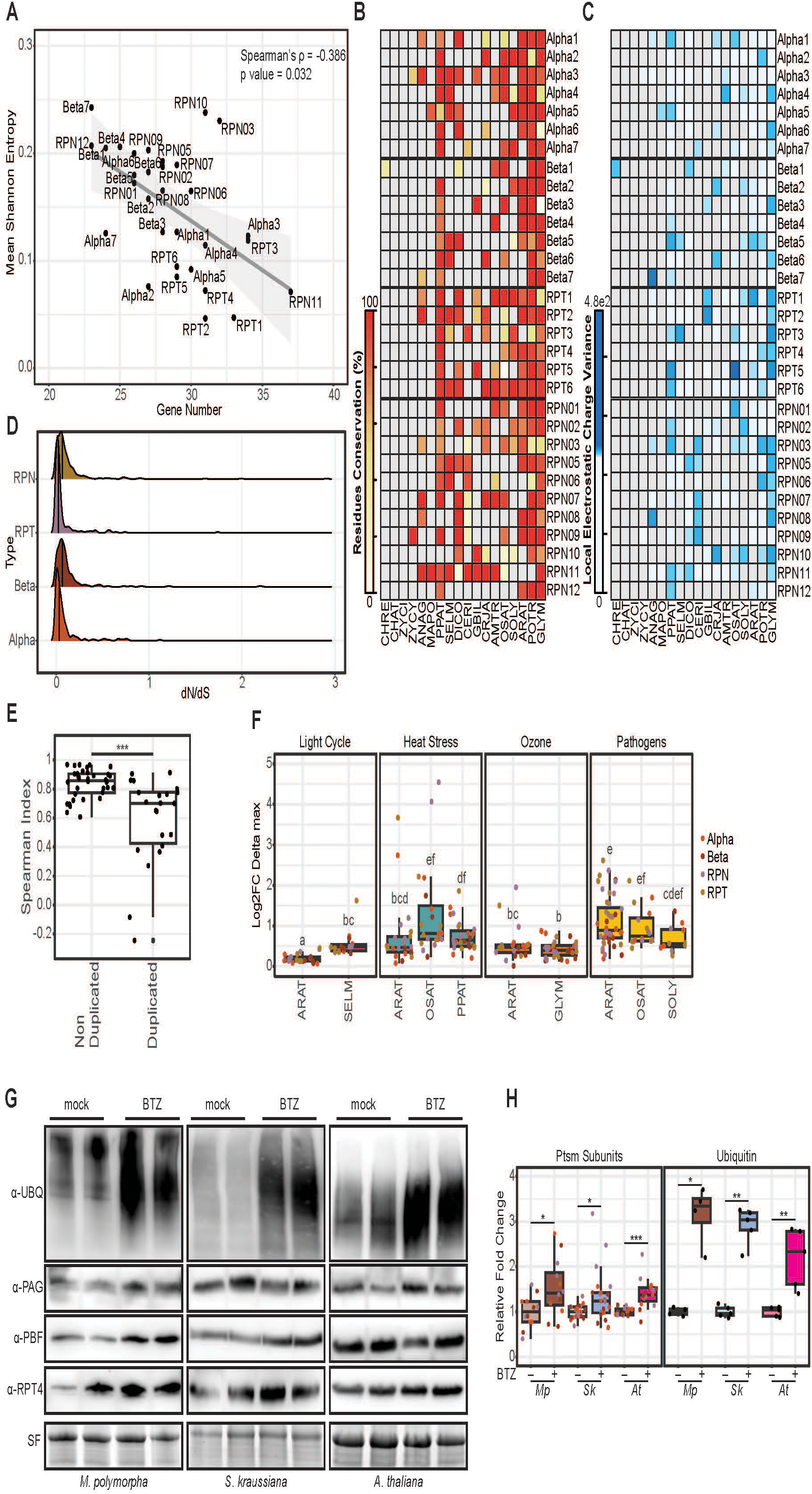
Proteasome paralogs diversified through sequence fine-tuning and divergent transcriptional regulation. **A.** Mean Shannon entropy (from multiple-sequence alignments) versus number of genes per proteasome subunit family; linear regression fit shown in gray. Spearman’s rank correlation rho and associated p value are indicated in the upper right corner. **B.** Conservation of post-translationally modified residues between paralogs within each species. **C.** Mean local electrostatic charge variance between paralogs within each species. **B-C**. Grey boxes indicate absence of gene duplication. **D.** Density plot of post-duplication dN/dS estimates for each subunit class. **E.** Transcriptional co-regulation in *A. thaliana* assessed by Spearman correlation between non-duplicated genes or between paralogs. Statistical significance by Wilcoxon rank-sum test (*** p < 0.001). **F.** Divergence in transcriptional regulation between paralogs across species and conditions. Lowercase letters indicate statistical groups determined by pairwise Wilcoxon rank-sum tests with false discovery rate correction (significant at q < 0.05). **E-F**. Correlations and divergences are based on Log2 FC data shown in Fig S11. **G.** Immunoblots for multiple 26S proteasome subunits and ubiquitinated proteins from crude extracts of *M. polymorpha*, *S. kraussiana* or *A. thaliana* after mock treatment or Bortezomib (20 µM, 6 h). Stain-free gel images serve as the loading control. **H.** Fold change in abundance of 26S proteasome subunits and ubiquitinated proteins relative to mock; data points are colored by protein. Quantification represents a pool of 3 independent experiments. Statistical differences were assessed by Wilcoxon rank-sum test (* p < 0.05, ** p < 0.01, *** p < 0.001).

Together, these analyses indicate that plant proteasome subunits evolved under pervasive purifying selection, with diversification arising primarily through fine-tuning of surface electrostatics and PTM landscapes rather than through widespread amino acid replacements or adaptive shifts. This mode of change is consistent with functional conservation of the complex accompanied by species- and paralog-specific modulation.

### Proteasome gene repertoire expansion led to paralog differential transcriptional regulation

The multimeric nature of the proteasome and the prevalence of duplicates suggests that both – precise control of 26S proteasome gene dosage balance (Birchler and Veitia, 2010) and interactions among paralogs, prompting potential paralog interference (Kaltenegger and Ober, 2015) – are important constraints on proteasome. Given the pervasive purifying selection observed, extensive neo- or sub-functionalization from amino acid changes seems unlikely; instead, paralog sub-functionalization may have been driven by spatial or temporal segregation via differential transcriptional regulation.

One fate of duplicated genes is pseudogenization through reduced transcriptional activity (Yang et al., 2011). To test this, we analyzed mRNA levels of proteasome duplicates in several species that exhibit pervasive duplication (**Fig S10**). Most genes showed substantial expression, indicating few cases of pseudogenization. We then compared median transcript levels of proteasome genes across representative tissues in multiple species. While some paralogs displayed transcriptional divergence, reflected by negative correlations in expression levels (**Fig S11**), most showed similar patterns across tissues/developmental stages.

Developmental context alone thus revealed limited divergence. However, *A. thaliana* harbors a proteasome stress regulon with paralog-biased activation during stress (Gladman et al., 2016), and proteasome transcriptional control is key to proteotoxic stress responses (Langin et al., 2026). With this in mind, we reanalyzed our transcriptomic datasets with respect to duplicates and found a clear bias in the regulation of *A. thaliana* paralogs under chemical and bacterial stress, with most duplicated genes showing stronger transcriptional response of one paralog (**Fig S12A**). To test whether such bias extends to other plants, we examined tomato and rice transcriptomes during pathogenic infection (Pombo et al., 2014; Hu et al., 2023) and observed similar paralog bias. Because proteasome transcriptional dynamics also occur under abiotic stress (Langin et al., 2026), we surveyed contexts linked to differential proteasome expression and identified ozone and heat treatments (Waldeck et al., 2017; Albihlal et al., 2018; Yu et al., 2019; Elzanati et al., 2020; Morales et al., 2021; Yang et al., 2023; Chang et al., 2024). In these conditions also, differential expressions preferentially affect certain paralogs (**Fig S12B, C**). Notably, we also detected paralog bias in the bryophyte *P. paten*s, suggesting that stress-responsive proteasome duplicates are broadly conserved in plants. We next examined circadian modulation. In *A. thaliana*, proteasome gene mRNA levels varied significantly across the day/night cycle with a paralog bias similar to that seen under stress (**Fig S12D**). Analysis of *S. moellendorfii* transcriptome under comparable conditions confirmed that proteasome transcriptional regulation is also a feature of earlier-diverging plants (**Fig S12D**). This analysis highlights a recurrent transcriptional response of 26S proteasome genes across several species and environmental conditions, characterized by paralog-specific bias. This suggests that some paralogs might act as general stress-responsive genes. To address this possibility, we compared the ranking of *A. thaliana* genes based on their transcriptional responses across the four datasets analyzed (**Fig. S12E**). This visualization revealed large differences in the distribution of genes from the least to the most responsive across the four conditions, with no obvious pattern of consistently stress-responsive genes. This indicates strong plasticity and specificity in differential paralog regulation.

To quantify these differences, we compared correlations in fold change between duplicated subunits and those among non-duplicated subunits in *A. thaliana* (**Fig 2E**). In this analysis, we hypothesized that non-duplicated genes should show strong co-expression given their unique presence in the genome whereas duplicated gene co-expression should be lower given differential transcriptional regulation between paralogs. Indeed, we found that non-duplicated subunits exhibited strong positive correlations, whereas duplicated subunits showed significantly lower correlations. We then assessed the magnitude of these differences across species and conditions. Across all species examined, differences in expression fold-change between duplicates were of similar magnitude and were primarily context-dependent (**Fig 2F**).

Thus, differential transcriptional regulation of proteasome duplicates is not restricted to *A. thaliana* and extends beyond flowering plants.

Our transcriptomic analysis revealed sub-functionalization of paralogs through differential transcriptional regulation while highlighting regulation of proteasome homeostasis through gene expression modulation in Bryophyte plant models. Proteasome homeostasis regulation in *A. thaliana* is characterized by accumulation of proteasome subunits upon proteasomal stress (Langin et al., 2026). To ask whether this response is conserved broadly across plants, we compared proteasome subunit abundance in *M. polymorpha*, *S. kraussiana*, and *A. thaliana* following chemical proteasome inhibition with BTZ. Immunoblot analyses showed subunit accumulation in bryophytes and lycophytes similar to *A. thaliana* (**Fig 2G, H**), suggesting that the proteasome regulatory feedback loop is conserved across plants.

Together, these results indicate that transcriptional regulation of proteasome is a key response to environmental perturbations across the green lineage. We found that sub-functionalization of 26S proteasome paralogs is driven by transcriptional segregation more so than amino acid sequence evolution. Furthermore, we highlight a common regulatory logic in plants, linking contextual fine-tuning of 26S proteasome homeostasis with species-specific gene repertoires. Importantly, these results underscore the central role of transcriptional regulation in the evolution of plant 26S proteasome regulation.

### Proteasome-associated cis-elements are highly diversified in plants

We found that the 26S proteasome complex exhibits conserved dynamic transcriptional regulation across diverse contexts and species. Given the conserved logic of proteasome homeostasis maintenance across kingdoms, this suggests a shared regulatory mechanism across plants; however, the only characterized mechanism to date is in *A. thaliana* (Gladman et al., 2016; Langin et al., 2026).

To further analyze this conserved regulatory regimen, we examined conservation of the proteasome associated cis-element across the green lineage. The PRCE in *A. thaliana* and PACE in *S. cerevisiae* are specifically enriched in proteasome promoters and show positional bias (**Fig S1**). Thus, we searched plant promoters for analogous enrichment patterns by mapping short DNA motifs in regions near the TSS (-400/+100) of proteasome genes versus the same regions of all genes or versus the upstream 1,600 bp (**Fig 3A**). This yielded genome-wide and positional enrichment scores. We retained the top 10 motifs per species by combined score (**Table S4**) and found low ambiguity motifs in 12 of 19 species. We refined motifs by analyzing flanking regions (±5 bp) in proteasome promoters and generated logos representing proteasome-associated cis-elements (**Fig 3A, B**). Notably, enrichment of an RGCCCA motif associated with the PRCE was observed in all five sampled mesangiosperms, but not in the basal angiosperm *A. trichopoda* (**Fig 3B**). This suggests that PRCE-mediated transcriptional regulation emerged shortly after the evolution of flowers.

**Figure 3.**
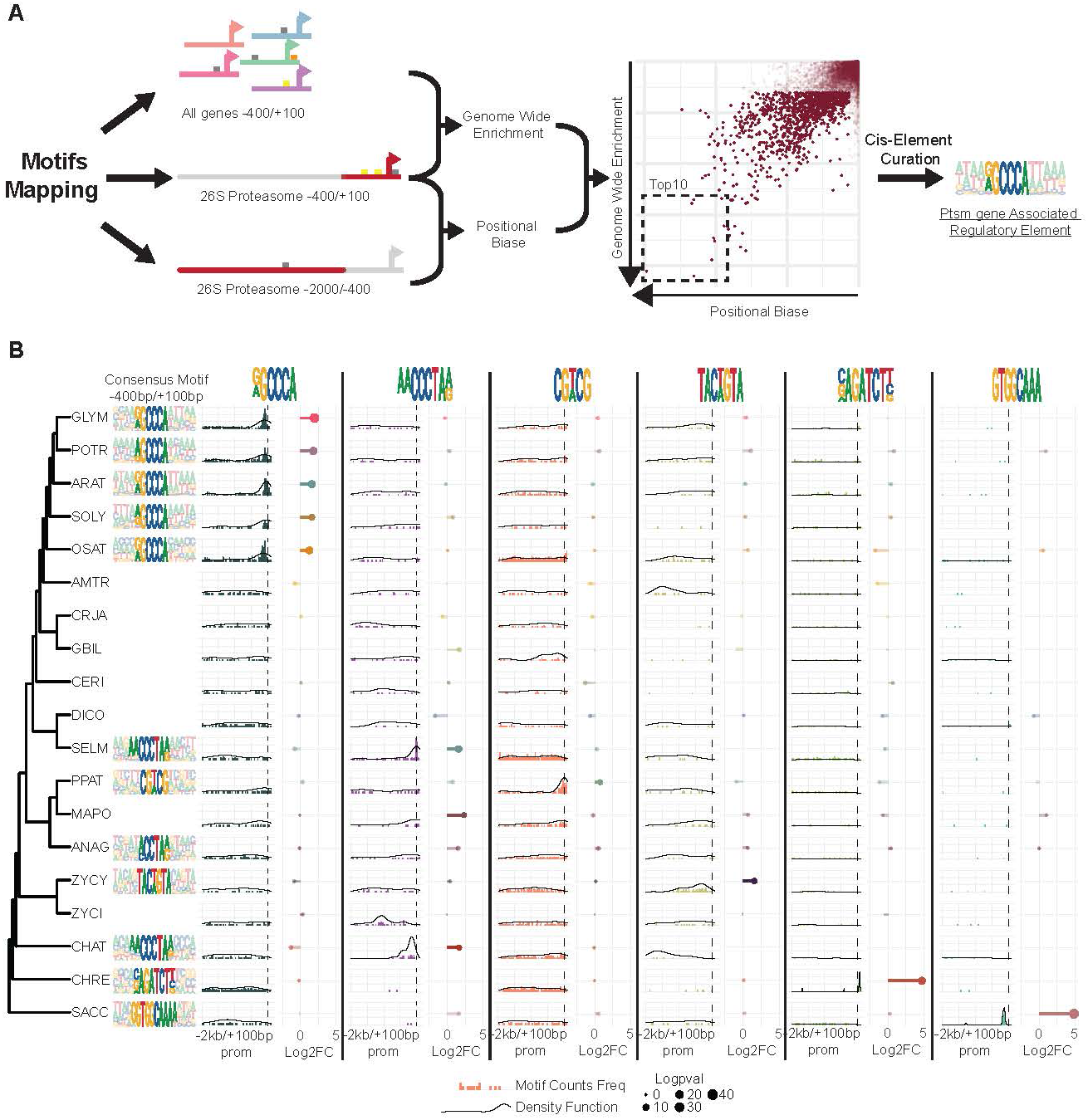
Proteasome associated cis-elements have largely diversified during plants evolution. **A.** Schematic of promoter DNA motif enrichment analysis in plant proteasome gene promoters. Short motifs were mapped in TSS-proximal regions of 26S proteasome genes and compared either to their frequency in all gene promoters or to distal regions within the same proteasome promoters. The resulting scores were combined to rank motifs; the top 10 candidates were examined, and a proteasome-associated cis-element was manually curated from them. **B.** Proteasome-associated cis-element analysis. For each species with a notable motif, consensus motif logos (promoter window -400 bp/+100 bp) are shown. For the six top candidate motifs, bar plots depict positional frequency along proteasome promoters (-2 kb/+100 bp); dotted lines mark the TSS. Black curves represent density functions. Lollipop charts display genome-wide enrichment as log2 fold change; lollipop size indicates -log10(p). Fisher’s exact test was used; lollipops are semi-transparent where p > 0.001.

To corroborate this, we assessed the evolutionary origin of the *A. thaliana* regulators NAC53 and NAC78. Phylogenetic analysis of NAC53/78 with other related NACs, AtNAC019, and their closest relatives from the six angiosperms and *M. polymorpha* showed that NAC53/78 form a distinct clade (**Fig S13A, B**). Consistent with the absence of PRCE in *A. trichopoda* proteasome promoters, no *A. trichopoda* proteins grouped within the NAC53/78 clade (**Fig S13A**), suggesting that the NAC53/78–PRCE module is specific to mesangiosperms. To validate conservation within mesangiosperms, we cloned NAC53/78 homologs from *S. lycopersicum* (SlNACs) and generated DNA probes from *S. lycopersicum* proteasome promoters bearing PRCEs (**Fig S13B**). As predicted, AtNAC53/78 and SlNACs specifically bound PRCE elements from *S. lycopersicum* promoters (**Fig S13B, C**). In parallel, both SlNACs – but not the distantly related MpNAC4– activated the *A. thaliana* RPT1a promoter to levels comparable to NAC78 (**Fig S13D**). These data validate conservation of the regulatory principles underlying 26S proteasome transcriptional regulation between *A. thaliana* and *S. lycopersicum* and strongly support broader conservation among flowering plants.

Outside angiosperms, we observed extensive diversification of proteasome-associated cis-elements, suggesting proteasome transcriptional regulatory modules have diversified across lineages. Notably, we found specific enrichment for the palindromic GCWGC, TACWGTA and VAGATCTB motifs in the distantly related species, *P. patens*, *Z. cylindricum* and *C. reinhardtii*, respectively. We also identified three species with related CCCTAR-containing motifs, two of which (*S. moellendorfii* and *C. atmophyticus*) shared a similar AACCCTAR motif (**Fig 3B**). To test interspecies conservation, we evaluated positional and genome-wide enrichment of each identified motif across all species. Strikingly, motifs were nearly always enriched only in the species in which they were identified (**Fig 3B**), confirming a high degree of diversification of proteasome-associated cis-elements.

To test whether proteasome-associated cis-elements underlie regulatory divergence among duplicates, we quantified motif frequencies in promoters of proteasome genes (-2kb/+100bp) (**Fig S14**). We frequently observed differences in motif counts between promoters of duplicated genes across all species examined (**Fig S14A, B**). Most pairs differed by one or two motifs, but extreme cases showed differences of up to eight motifs between duplicates (**Fig S14C**). These suggest that paralogs promoters could diversify through differential enrichment of proteasome-associated cis-elements.

Together, these analyses confirm conservation of the angiosperm proteasome regulatory module while suggesting that the broadly conserved feedback loop operates through highly species-specific cis-regulatory codes across plant evolution. Although proteasome gene duplications are generally recent, their promoters have diversified markedly between duplicates. This illustrates exceptional evolutionary plasticity in the regulation of one of the most conserved molecular complexes in eukaryotes.

### Telomere repeat binding factors are central to proteasome transcriptional regulation in plants

Our analysis of proteasome-associated cis-elements identified CCCTAR as the predominant motif of proteasome promoters in three non-seed lineages. This observation suggests two hypotheses: 1) an ancient role of the motif in proteasome transcriptional regulation or, 2) convergent evolution fixing the motif in 26S proteasome promoters from several plant lineages. The motif corresponds to a telobox motif, related to telomeric repeats (Bilaud et al., 1996), which is recognized by telomere repeat-binding factors (TRBs) in plants, resulting in recruitment of the Polycomb repressive complex to regulate chromatin accessibility (Zhou et al., 2018; Amiard et al., 2024). ChIP-seq studies further show that TRBs bind not only telobox but also the site II RGCCCA motifs (Zhou et al., 2016; Wang et al., 2023; Amiard et al., 2024), which correspond to the angiosperm PRCE (**Fig 3B**). These observations suggest that TRBs could recognize proteasome-associated cis-elements in at least eight of the species we analyzed, including *A. thaliana*.

To test whether TRBs bind the PRCE, we purified *A. thaliana* TRB1, TRB3, and TRB5 and performed EMSAs using probes containing the PRCE. We found that TRBs bound PRCE motifs *in vitro* with a partial specificity toward the TGGGC/GCCCA core sequence (**Fig 4A, B**), supporting TRB recognition of PRCE. We next assessed *in vivo* association by reanalyzing published AtTRB ChIP-seq datasets around 26S proteasome gene TSSs. All four TRBs examined showed frequent enrichment near TSSs (**Fig S15A**). Because TRBs primarily act through local chromatin remodeling, we examined chromatin status of 26S proteasome promoters during bacterial infection (Ding et al., 2021); a context inducing differential regulation of 26S proteasome genes (Langin et al., 2026). Several 26S proteasome promoters displayed significant chromatin opening near the TSS (**Fig S15B**). TRB ChIP-seq signals were specifically enriched in proteasome promoters’ regions harboring a PRCE or showing increased accessibility during infection (**Fig S15C**). Together, these data indicate that TRB proteins associate with 26S proteasome promoters *in vivo* via PRCE-containing regions and support that TRBs could act as novel candidate regulators of 26S proteasome transcription.

**Figure 4.**
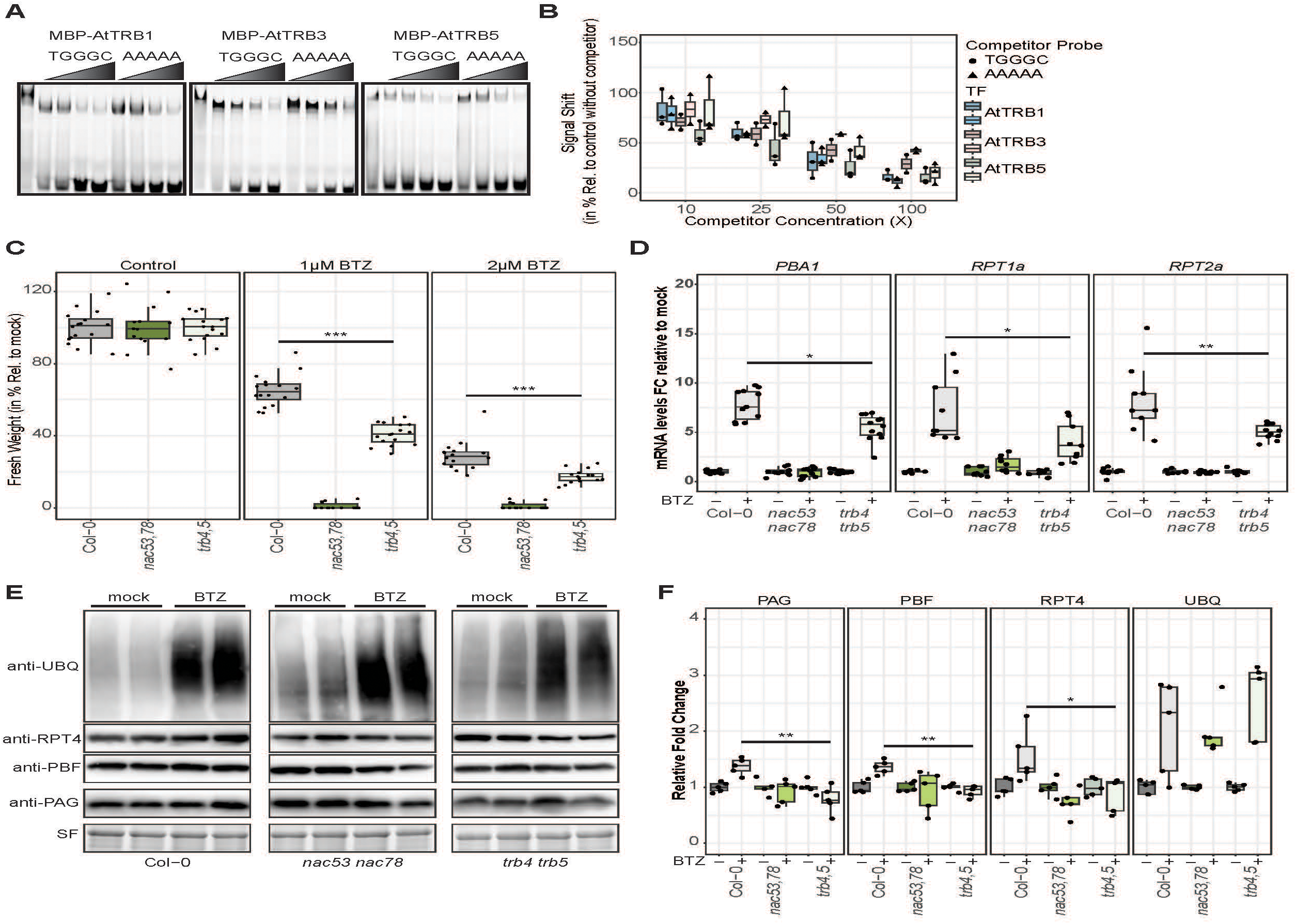
TRB4 and TRB5 regulate 26S proteasome homeostasis in *A. thaliana*. **A.** Electrophoretic mobility shift assay (EMSA) competition with MBP-tagged AtTRB1/3/5 using SlPBF probe from Fig S12. Binding-induced shifts in the first lanes were challenged with increasing concentrations (10× to 100×) of unlabeled competitor probes: wild type (TGGGC) or mutated (AAAAA). **B.** Quantification of shifted signal expressed as percentage of the no-competitor control shift for each competitor’s concentration. Box colors denote MBP-AtTRB proteins; point shapes denote competitor type. n = 3 independent experiments for AtTRB1 and AtTRB5; n = 2 for AtTRB3. **C.** Seedling fresh weight at 10 days after germination under the indicated treatments, expressed as percent of mock. Statistical significance by Wilcoxon rank-sum test (*** p < 0.001). **D.** mRNA level Fold Change of the indicated transcripts in Col-0, *nac53-1 78-1* or *trb4,5* seedlings (7 days old) after mock or Bortezomib (20 µM, 6 h) treatment; fold change relative to mock. Statistical significance by Wilcoxon rank-sum test (* p < 0.05, ** p < 0.01). **E.** Immunoblots for multiple 26S proteasome subunits and ubiquitinated proteins from crude extracts of Col-0, *nac53-1 78-1* or *trb4,5* seedlings (7 days old) after mock or Bortezomib (20 µM, 6 h). Stain-free gel image serves as loading control. **F.** Fold change in abundance of 26S proteasome subunits and ubiquitinated proteins relative to mock. Quantification from 3 independent experiments. Statistical significance by Wilcoxon rank-sum test (* p < 0.05, ** p < 0.01).

To validate TRB function, we analyzed *trb* mutants under proteasome inhibition. Based on prior evidence that AtTRB4 and AtTRB5 act as transcriptional activators (Amiard et al., 2024), we focused on these paralogs. We found that mutant plants *trb4 trb5* were hypersensitive to chemical proteasome inhibition (**Fig 4C**). Consistent with this phenotype, induction of 26S proteasome transcripts and accumulation of 26S proteasome subunits were significantly impaired in *trb4,5* mutants (**Fig 4 D,E,F**). These results support a role for TRB4 and TRB5 in activating 26S proteasome gene expression in *A. thaliana*.

In sum, despite extensive diversification of proteasome cis-elements, many of the predominant cis-elements across species correspond to known TRB targets, including the PRCE in angiosperms. We validate TRBs as positive regulators of proteasome genes in *A. thaliana* through the isoforms TRB4/5. The origin of TRB-targeted cis-elements in plant proteasome promoters suggests an ancestral role for TRBs in proteasome transcriptional regulation. This conserved role, coupled with diversification of proteasome-associated cis-elements, suggests that evolution of the proteasome regulatory module is linked to other regulatory processes.

### Plant 26S proteasome is embedded in a gene regulatory network

We recently showed that proteasome transcriptional activators in *A. thaliana* coordinately regulate 26S proteasome genes and photosynthesis-associated genes via the PRCE (Langin et al., 2026). Similar transcriptional coupling between 26S proteasome genes and other pathways has been reported in other model organisms (Krämer et al., 2021; Ruvkun and Lehrbach, 2023). TRBs are also implicated in regulating multiple processes (Zhou et al., 2016; Amiard et al., 2024). These observations suggest that proteasome activators not only maintain 26S proteasome homeostasis but also embed it into transcriptional networks. Moreover, the diversification of proteasome gene repertoires and of proteasome-associated cis-elements occurred on different scales, implying that changes in cis-regulatory logic are not solely driven by proteasome gene evolution. We therefore hypothesized that proteasome-associated cis-elements link the 26S proteasome to a wider network of co-functional proteins that is maintained throughout plant evolution.

To test this, we constructed a protein co-evolution network centered on plant 26S proteasome subunits using Evolutionary Rate Covariation (ERC) analyses, implemented in the ERCnet algorithm (Forsythe et al., 2025). Previous ERC analyses revealed strong signatures of ERC within and between plastid proteostasis pathways, revealing connectivity between co-functional pathways within a larger interactome network (Gatts et al., 2026). Our 26S proteasome-focused network revealed that, while pairwise combinations of proteasome subunits display elevated ERC signature compared to the genome-wide background (**Fig S16**), only 7 out of the 465 possible pairs were deemed significant ERC hits (**Table S5**). The moderate level of ERC among proteasome subunits supports largely independent sequence evolution among subunits in plants. However, we identified 386 non-proteasome gene families with significant ERC to at least one proteasome subunit gene family (**Table S5**), defining a conserved proteasome-associated protein network (**Fig 5A**). Network members showed numerous significant ERC relationships to other non-proteasome members, and clustering revealed distinct modules, indicating that proteasome subunits co-evolved within a network comprising both proteasome-dependent and -independent relationships. Gene Ontology analysis (based on *A. thaliana* orthologs) highlighted enrichment for trafficking, RNA splicing, chromatin and chloroplast components (**Fig S17A, B**), embedding the 26S proteasome within a broader proteostasis framework.

**Figure 5.**
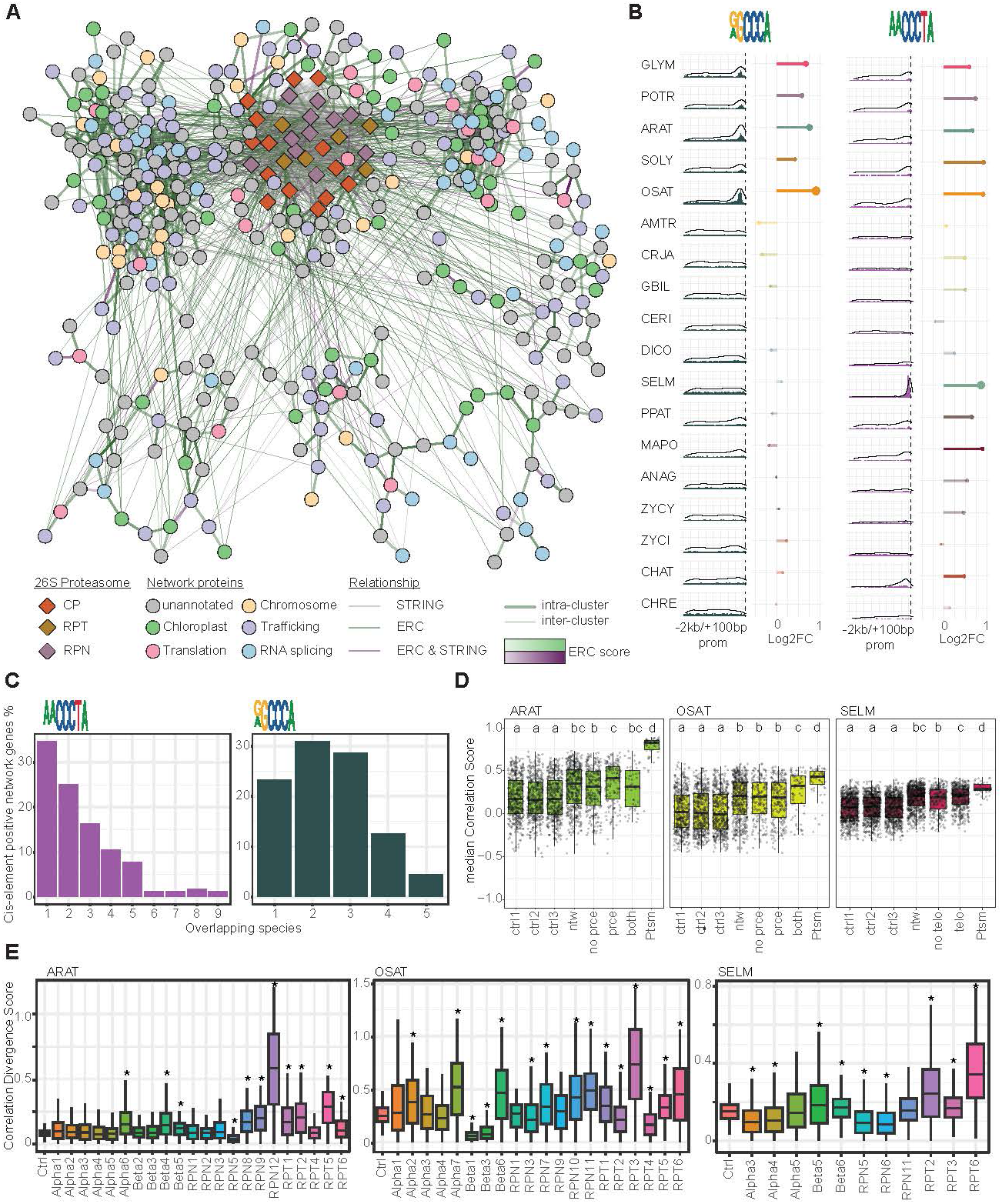
Proteasome regulation is embedded within a proteostasis-centered regulatory network. **A.** Evolutionary rate covariation (ERC) network centered on 26S proteasome subunits. Proteasome nodes are diamonds colored by subunit class; other proteins are circles colored by enriched Gene Ontology (GO) terms (see Fig S15). Edges indicate associations from STRINGdb, ERC, or both (color-coded); edge thickness distinguishes intra- versus inter-cluster links; edge transparency reflects ERC score magnitude. **B.** Enrichment and positional analyses of [GA]GCCCA (PRCE/site II-like) and AACCCTA (telobox) motifs in promoters (-2 kb/+100 bp) of network genes across species. Bar plots show motif positional frequency; dotted lines mark the TSS. Black curves represent density functions. Lollipop charts display genome-wide enrichment as log2 fold change; lollipop size indicates -log10(p). Fisher’s exact test was used; lollipops are semi-transparent where p > 0.001. **C.** Proportion of overlapping genes bearing proteasome-associated cis-elements in the proximal promoter (-400 bp/+100 bp) across species showing significant enrichment for the respective motifs. **D.** Median Spearman correlation of gene expression with proteasome genes for each species. Controls are three sets of randomly sampled genes. Groups: “ntw” (all network genes from panel A); “no prce” (network genes lacking [GA]GCCCA in -400/+100); “prce” (with [GA]GCCCA); “no telo” (lacking AACCCTA); “telo” (with AACCCTA); “both” (with both motifs); “ptsm” (proteasome genes). Statistical groups were assessed by pairwise Wilcoxon rank-sum tests with FDR correction (q < 0.05). **E.** Divergence in co-regulation of proteasome paralogs with network genes, shown as the delta in Spearman correlation between paralogs. Statistical differences versus the control group were assessed by pairwise Wilcoxon rank-sum tests with FDR correction (* q < 0.05).

We next asked whether this co-evolution network is connected through shared transcriptional regulation. Using MEME Suite, we searched for enriched promoter elements among network members, excluding 26S proteasome genes (Bailey et al., 2015). We detected enrichment of TGGGC- or AACCCT-related motifs across multiple species (**Fig S17C**). TGGGC-related motifs appeared only in angiosperms, mirroring their enrichment in proteasome promoters, whereas telobox-related motifs (AACCCT) were found in several diverging species including *A. atmophyticus* and *S. moellondorfii*, paralleling our proteasome promoter analyses (**Fig 3B**). Detailed enrichment tests confirmed specific enrichment of the RGCCCA motif in angiosperms and of the AACCCTA telobox motif in nine species (**Fig 5B**). Consistently, analysis of the two additional streptophyte proteasome-associated cis-elements revealed positional and genome-wide enrichment in species (or close relative) where the motif is enriched in proteasome promoters (**Fig S17D**). These results indicate that plant 26S proteasome evolved within a proteostasis-related protein network linked by proteasome associated cis-elements diversification.

We then assessed conservation of the genes bearing RGCCCA or AACCCTA motifs across species by comparing overlaps of orthogroups containing the same cis-elements. For telobox-positive genes, nearly 35% were species-specific, and ∼75% were shared by no more than three of the nine species examined (**Fig 5C, Fig S17E**), indicating substantial diversification. PRCE-positive genes in angiosperms were also diversified: >75% overlapped in no more than three of the five species (**Fig 5C, FigS17F**). This suggests that transcriptional co-regulation between proteasome genes and network members has been repeatedly rewired across plant evolution.

To validate co-regulation, we tested whether network genes are co-expressed with proteasome genes. Indeed, network genes showed significantly stronger expression correlations with proteasome genes than background groups (**Fig 5D**). In *A. thaliana*, genes with PRCEs in the proximal promoter (−400/+100) were most strongly correlated (**Fig 5D**). In

*O. sativa*, genes harboring both PRCE and AACCCTA elements showed the strongest correlations (**Fig 5D**). In *S. moellendorfii*, network genes were more strongly co-regulated with proteasome genes overall, with AACCCTA-positive promoters showing significantly higher correlations (**Fig 5D**). These patterns support a link between proteasome-associated cis-elements in network gene promoters and their co-regulation with proteasome genes.

Finally, we asked whether proteasome duplications shape these transcriptional connections. To assess if paralogs are differentially connected to the network, we quantified the divergence between paralogs based on mRNA level correlations with network genes. A high divergence score indicates strong differences in how the duplicated genes are co-regulated with the network. As a control score, we used the median divergence calculated between non-duplicated genes. Duplications generally increased divergence (**Fig 5E**), indicating that differential regulation between duplicates is embedded within the broader network. However, some duplicates showed similar co-regulation with network genes, underscoring that each duplication yields a distinct regulatory outcome. These results demonstrate a transcriptional connection between the proteasome-associated network and proteasome duplicates, linking the two main drivers of 26S proteasome evolution in plants.

In summary, proteasome subunits have co-evolved with a proteostasis-centered protein network. Promoters of network genes are recurrently enriched for proteasome-associated cis-elements, yet the specific genes bearing these motifs differ among species, while remaining systematically co-regulated with the proteasome. We further show a direct connection between the transcriptional regulation of proteasome duplicates and the network. These findings demonstrate that evolution of plant proteasome regulation is embedded within a regulatory network driven by diversification and differential enrichment of associated cis-elements.

## Discussion

### Proteasome regulation in A. thaliana involves a complex transcriptional regulatory module

In this study we identified telomere repeat–binding proteins (TRBs) as transcriptional regulators of 26S proteasome genes in *A. thaliana*, binding the same proteasome-associated cis-element (PRCE/site II) recognized by NAC53/78 (Langin et al., 2026). This supports a coordinated mechanism in which TRBs and NAC53/78 jointly control proteasome homeostasis (**Fig 6A**). Consistent with known roles of TRBs in recruiting chromatin-remodeling complexes (Zhou et al., 2018; Wang et al., 2023; Amiard et al., 2024), *A. thaliana* proteasome promoters bound by TRBs show signatures of chromatin remodeling. We propose that TRB4/5 engage PRCE at proteasome promoters, recruit remodeling machinery, and increase accessibility for NAC53/78, which then activate transcription (**Fig 6A**). Delineating the temporal interplay between TRBs and NAC53/78 at proteasome promoter and the significance of chromatin remodeling on subsequent regulation would be essential to validate our model.

**Figure 6.**
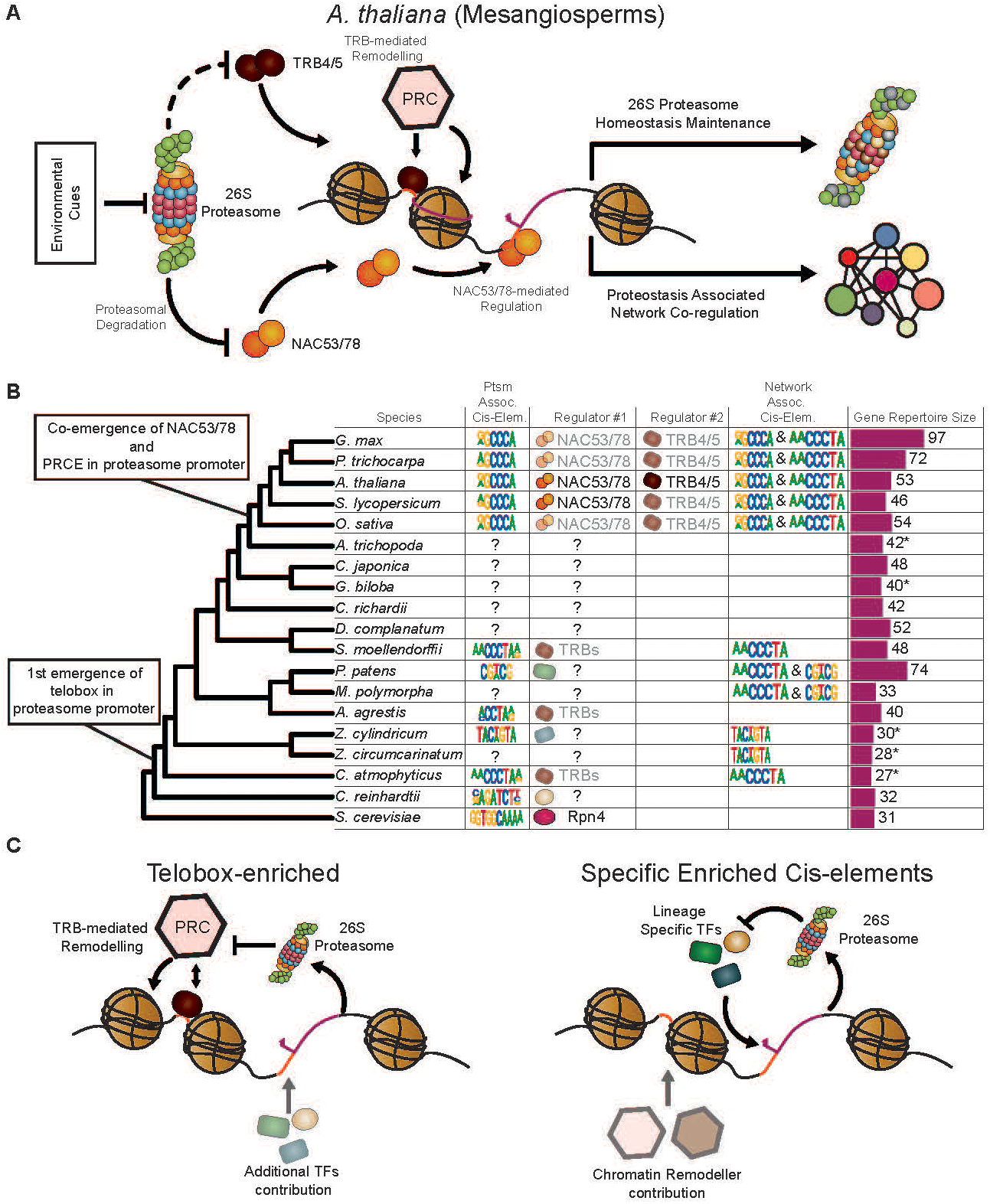
Working model of 26S Proteasome Homeostasis regulation and associated evolutionary features in plants. **A.** Proposed model of 26S proteasome transcriptional regulation in A. thaliana. Environmental cues reduce 26S proteasome activity or capacity, triggering a feedback response mediated by NAC53/78 and TRB4/5. TRB4/5 associate with proteasome promoters and promote chromatin remodeling, potentially through recruitment of Polycomb repressive complex (PRC)-associated remodeling activity, thereby facilitating NAC53/78-mediated transcriptional regulation through PRCE/site II. This regulatory module contributes to maintenance of 26S proteasome homeostasis and co-regulation of a broader proteostasis-associated network. Dashed inhibitory links indicate hypothetical proteasome-dependent turnover of TRB4/5. **B.** Evolutionary model of proteasome-associated cis-regulatory mechanisms across the lineages. The framework summarizes enriched cis-elements in 26S proteasome promoters, candidate or validated regulators, enriched cis-elements in proteasome-associated network genes, and proteasome gene repertoire size. The first enrichment of telobox-like motifs in proteasome promoters is inferred near the base of streptophytes, whereas co-emergence of NAC53/78 and PRCE enrichment in proteasome promoters is inferred in mesangiosperms. Transparency indicates speculative regulators. Question marks indicate unknown or unresolved regulators/cis-elements. Asterisks indicate species for which proteasome gene repertoire size may be underestimated because of incomplete annotation or missing subunits. **C.** Proposed scenarios for diversification of proteasome regulatory mechanisms. In telobox-enriched lineages, TRBs may contribute primarly to proteasome homeostasis regulation, potentially together with additional transcription factors. In lineages enriched for specific uncharacterized cis-elements, distinct transcription factors may bind these motifs and cooperate with chromatin remodelers to regulate proteasome genes. These scenarios illustrate how conserved regulatory logic may be maintained while cis-elements and transcriptional regulators diversify across plant evolution.

The proteasome regulatory feedback loop relies on constitutive turnover of the transcriptional activators (Wang et al., 2010; Ruvkun and Lehrbach, 2023; Langin et al., 2026). By analogy, TRBs could also be subject to proteasomal degradation, although this remains to be tested in plants; proteasome-mediated turnover of telomere-binding proteins is documented in other eukaryotes (Lin et al., 2015; Zalzman et al., 2020). Investigating mechanisms of TRB turnover would permit us to determine if TRBs are integral components of the proteasome regulatory feedback loop

TRB1/2/3 have been described primarily as repressors, whereas TRB4/5 act as activators (Zhou et al., 2016; Zhou et al., 2018; Amiard et al., 2024). Notably, NAC53/78 can also mediate repression of photosynthesis-associated nuclear genes via PRCE (Langin et al., 2026), and this motif is enriched among chloroplast-related genes differentially expressed in *trb1-1* (Zhou et al., 2016). These observations suggest that TRBs and NAC53/78 cooperate beyond proteasome genes to coordinate a broader proteasome-associated transcriptional network (**Fig 6A**).

### Diversification of proteasome cis-regulatory mechanism highlights distinct evolutionary trajectories

Our investigation into the role of TRBs in 26S proteasome gene regulation in *A. thaliana* was motivated by the discovery of TRB-targeted cis-elements enriched in proteasome promoters across several non-angiosperm species, including streptophyte algae (*C. atmophyticus*), bryophytes (*A. agrestis*) and lycophytes (*S. moellendorfii*) (**Fig 6B**). In addition, telobox motifs appear enriched in promoters of proteasome-associated network members in half of the surveyed species, including the other two bryophytes examined (*M. polymorpha* and *P. patens*). These patterns motivate two hypotheses: i) recurrent evolutionary convergence fixing TRB involvement in proteasome and/or associated network regulation, or ii) an ancestral, TRB-dependent mode of proteasome regulation which emerged in streptophytes. The conserved motif enrichment across deeply diverged lineages, the broad conservation of TRBs in streptophytes (Kusová et al., 2025), and the tight functional link between telobox and PRCE/site II (Trémousaygue et al., 2003; Gaspin et al., 2010) collectively favor the ancestral-trait hypothesis.

In this scenario, TRB-dependent proteasome regulation emerged alongside TRB evolution in streptophytes through the enrichment of telobox motifs in 26S proteasome promoters (Kusová et al., 2025) and was subsequently maintained to varying degrees across lineages (**Fig 6B**). In parallel, novel cis-elements appear to have arisen in proteasome promoters, including the palindromic GCWGC in *P. patens,* TACWGTA in *Z. cylindricum* and the PRCE in angiosperms. PRCE shares notable features with the telobox: it is bound by TRBs, often occurs near teloboxes in angiosperm promoters (Trémousaygue et al., 2003; Gaspin et al., 2010) and contains a C-rich core. We therefore propose that PRCE in angiosperm proteasome and associated-network promoters may have evolved from telobox-like precursors, retaining TRB recognition while accommodating regulation by the NAC53/78 clade (**Fig 6B**). Further characterization of this scenario through extensive analysis across phylogeny would provide a powerful case study for *de novo* cis-regulatory sequence emergence.

Despite an apparently conserved regulatory logic, proteasome-associated cis-elements are diverse and often lineage specific. Several non-mutually exclusive scenarios could explain this pattern. In telobox-bearing species, proteasome homeostasis may rely primarily on TRB function to activate proteasome transcription potentially with additional, specialized transcription factors that recognize telobox motifs and participate in the feedback loop alongside TRBs (**Fig 6C**). In species with lineage-specific proteasome-associated motifs, distinct transcription factors likely evolved to bind those elements, while TRBs or other conserved recruiters of chromatin remodelers may have adapted to cooperate at these sites (**Fig. 6C**). Functionally validating TRB-dependent proteasome regulation in divergent lineages and identifying the transcription factors that recognize lineage-specific motifs will clarify the origin and diversification of this regulatory system.

### Proteasome duplicates transcriptional divergence and sequence fine-tuning suggest different regulatory functions

A prominent feature of 26S proteasome evolution is expansion of its gene repertoire. Most paralogs in our dataset appear to be of recent origin, yielding lineage-specific gene sets. Ancient whole-genome duplications (WGDs) that shaped green lineage evolution (Tiley et al., 2016) contributed little to proteasome gene expansion, whereas more recent WGDs associated with lineage-specific innovations – for example in Brassicales, Glycine, and mosses (Lang et al., 2018; Mabry et al., 2020) – appear more influential.

Analyses of duplicate coding sequence evolution indicate only moderate functional divergence among 26S proteasome paralogs. By contrast, transcriptional divergence is substantial, consistent with the high fraction of duplicates showing expression differences in *A. thaliana* and *G. max* (Ganko et al., 2007; Roulin et al., 2013). Notably, whereas prior work emphasized tissue-specific divergence for gene duplicates, our data indicate that proteasome paralog expressions diverge primarily in response to environmental cues. This echoes the role of CP subunit duplication in animals in generating context-specific proteasomes (Murata et al., 2018). Plants may likewise deploy stress-responsive 26S proteasomes such as the animal immunoproteasome – an attractive possibility given the central role of the proteasome in plant immunity (Üstün et al., 2018; Langin et al., 2023; Langin et al., 2026). This hypothesis is further supported by the divergence of PTM sites between paralogs, given the strong influence of PTMs on 26S proteasome activity (Hirano et al., 2016; Kors et al., 2019). Furthermore, emerging evidence points that proteasome trafficking is a key regulatory axis (Albert et al., 2020; Zhang et al., 2023; Karimi et al., 2025; Tang et al., 2026). Preferential paralog expression could thus bias trafficking and assembly, adjusting spatiotemporal proteasome regulation under stress. These observations support a model in which plants evolve species-specific repertoires of proteasome genes to tailor proteasome regulation to environmental demands. Understanding the functional and regulatory role of these stress-responsive paralogs will be an important subject for future research.

### The diversity of transcriptional regulatory module connects proteasome within plant proteostasis networks

Pathways that regulate proteostasis are among the most conserved in eukaryotes, alongside fundamental mechanisms of metabolism and development (Cox et al., 2026). The existence of a coordinated proteostasis network is widely accepted (Balchin et al., 2016); however, studies mostly focus on individual functional nodes – such as Hsp chaperones or mTORC1 signaling (Taipale et al., 2014; Zhang et al., 2014) – with limited global views of the network.

Our analysis uncovered a protein network whose members have co-evolved with 26S proteasome subunits. Notably, these genes show signatures of co-evolution not only at the protein-sequence level but also through specific enrichment of proteasome-associated cis-elements. This supports a coordinated proteostasis network whose components are transcriptionally co-regulated with proteasome genes via shared cis-regulatory architecture. This network is enriched for proteins involved in canonical proteostasis mechanisms, including regulation of protein synthesis and trafficking. While there is growing evidence that evolutionary rate covariation indicates non-physical co-functional links between proteins (Clark et al., 2012; Forsythe et al., 2021; Little et al., 2024), there are ample examples of physically interacting proteins exhibiting tight co-evolution via compensatory evolution (Pazos and Valencia, 2008). This implies that proteasome subunits could serve as a physical interface with other multimeric complexes, such as the spliceosome or SNARE complexes. Investigating these relationships in more detail would be an important step toward understanding of the proteostasis network.

Interestingly, this network isn’t restricted to proteostasis mechanisms, as it also displays enrichment for chloroplast components and mitosis factors, indicating intimate connections to energy metabolism and the cell cycle. The former aligns with the characterization of NAC53/78 as coordinators of photosynthesis and proteasome homeostasis in *A. thaliana* (Langin et al., 2026). The latter connects to previous studies highlighting the importance of telobox and PRCE/site II motifs in regulating cell cycle–related genes (Trémousaygue et al., 2003). These data indicate that the proteasome is embedded in a conserved gene regulatory network coordinating proteostasis with energy metabolism and the cell cycle. Investigating the conservation of this network outside plants and its specificity in distinct lineages would be important to better understand eukaryotic evolution.

Together, our results indicate that the plant 26S proteasome has evolved along two complementary axes: an expanding, lineage-specific gene repertoire, and an increasingly diversified cis-regulatory landscape built upon a conserved TRB-dependent regulatory core. This architecture allows a structurally and functionally conserved molecular machine to be flexibly integrated into the shifting regulatory demands of each plant lineage — and, more broadly, into the wider proteostasis network with which it co-evolves.

## Author contributions

S.U. and G.L. conceived the project. S.U. and G.L. supervised the experiments. G.L designed and performed the experiments. T.W. and D.B contributed to the experiments. E.S.F. and G.L. supervised the bioinformatic analysis. G.L designed and performed bioinformatic analysis. E.S.F. and E.A.R contributed to the bioinformatic analysis.

## Acknowledgment

We thank Dr. Simon Amiard from iGReD, CNRS and Dr. Franziska Turck from MPI for Plant Breeding Research, for sharing plant lines and plasmids concerning TRBs. We thank Dr. Hee Sung Fiona Kang from Ruhr-University Bochum for sharing *M. polymorpha* plants. We thank the RUB Botanical Garden for sharing *S. kraussiana* plants.

## Data availability statement

The scripts used for the analysis and the resulting data are available upon request from the corresponding authors.

## Supplemental Data

**Figure S1.**
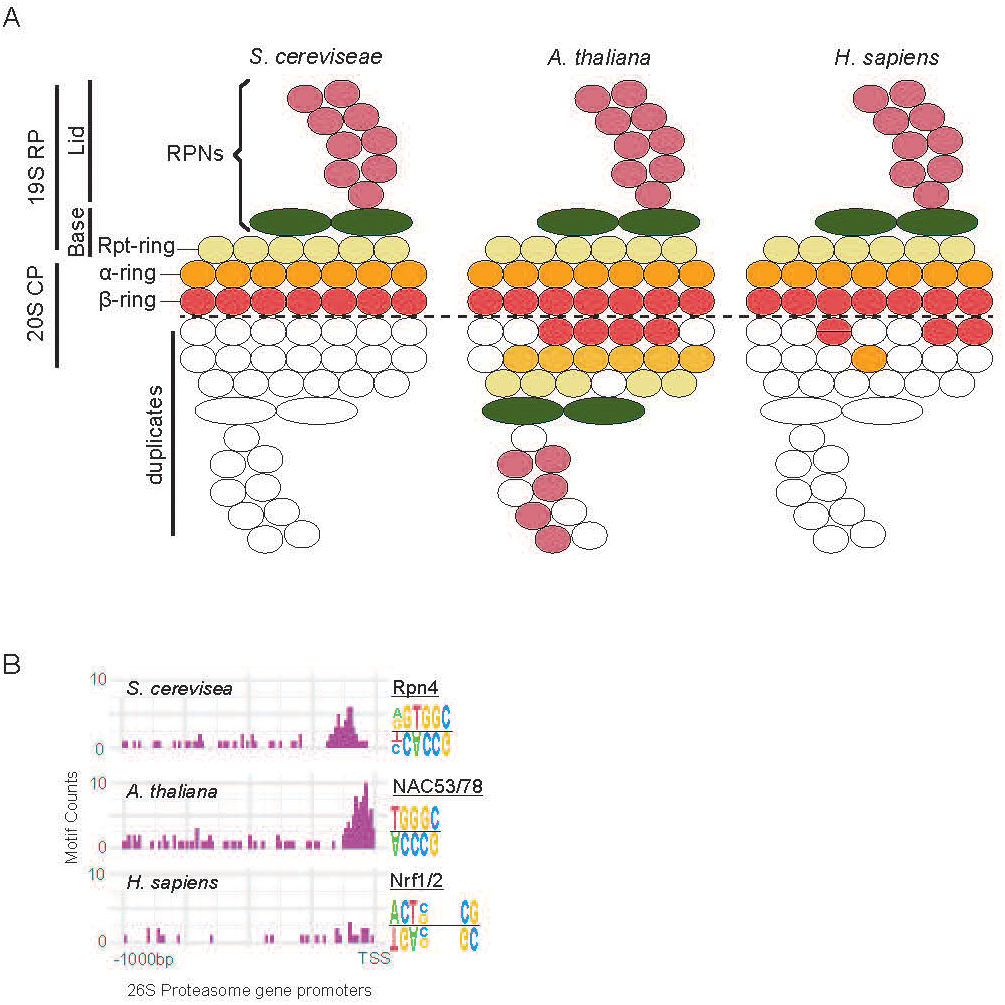
Comparison of 26S proteasome features in three model eukaryotes. **A.** Size of the 26S proteasome gene repertoire in *S. cerevisiae*, *A. thaliana* and *H. sapiens*. Duplicated subunits are indicated by colored circles in the lower half of the 26S proteasome schematic. **B.** Known proteasome-associated cis-elements: motif logos, counts, and positional distributions within proteasome promoters (-1 kb from TSS). Names of transcription factors reported to bind these elements in proteasome promoters are indicated.

**Figure S2.**
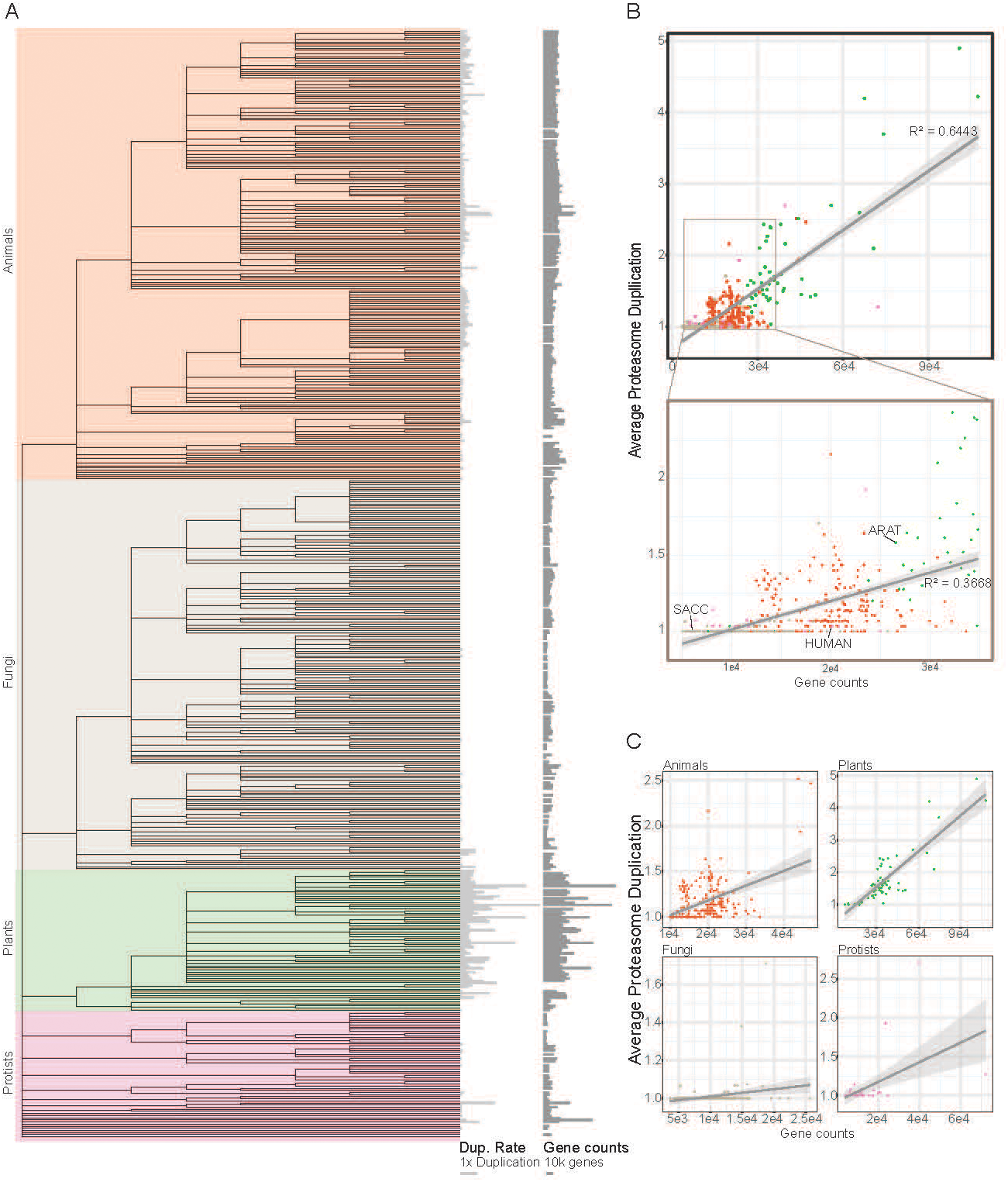
Large-scale analysis of proteasome gene duplication rate as a function of genome gene number. **A.** Phylogenetic tree of species extracted from the OMA database; colors denote major eukaryotic kingdoms. For each species, average proteasome duplication rate and total gene number are indicated to the right of the tree. **B.** Average proteasome duplication rate versus total gene number across species; inset zooms on species with 5,000–35,000 genes. In both plots, a linear regression fit is shown in gray with the corresponding R². **C.** Average proteasome duplication rate versus total gene number stratified by kingdom, with linear regression fits shown in gray.

**Figure S3.**
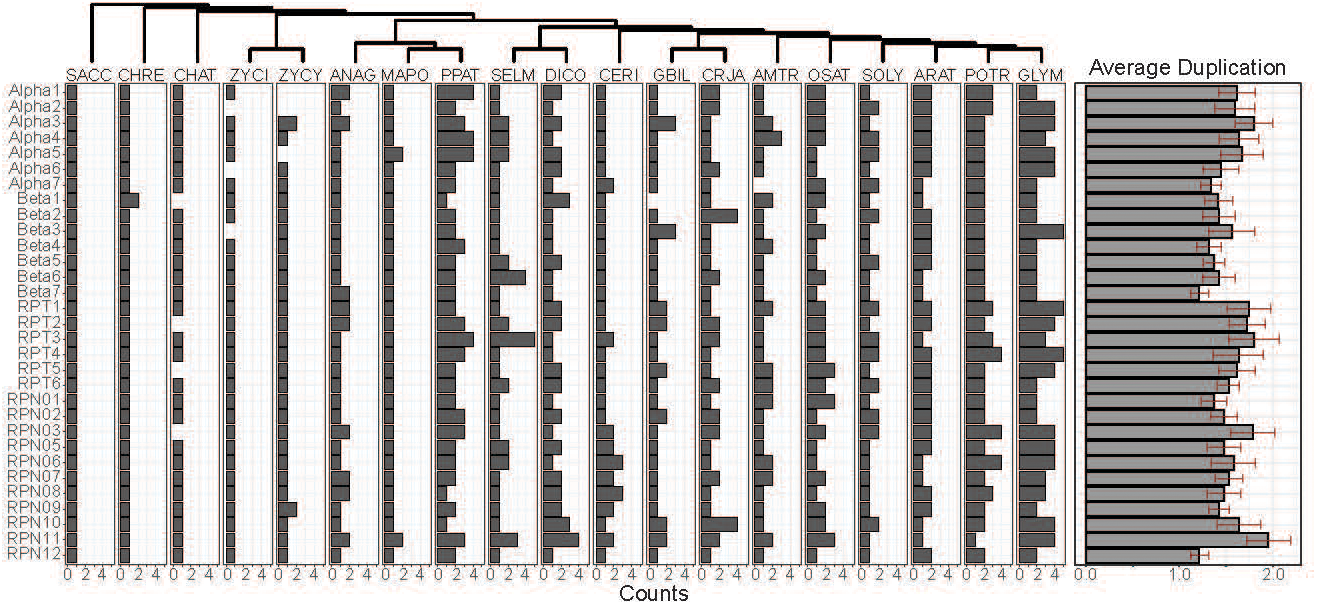
Detailed view of proteasome gene duplication across representative plant species. Counts of genes identified for each 26S proteasome subunit family are shown for each representative plant species. On the right, a bar plot represents the average duplication level per subunit family across plants; error bars denote standard deviation.

**Figure S4.**
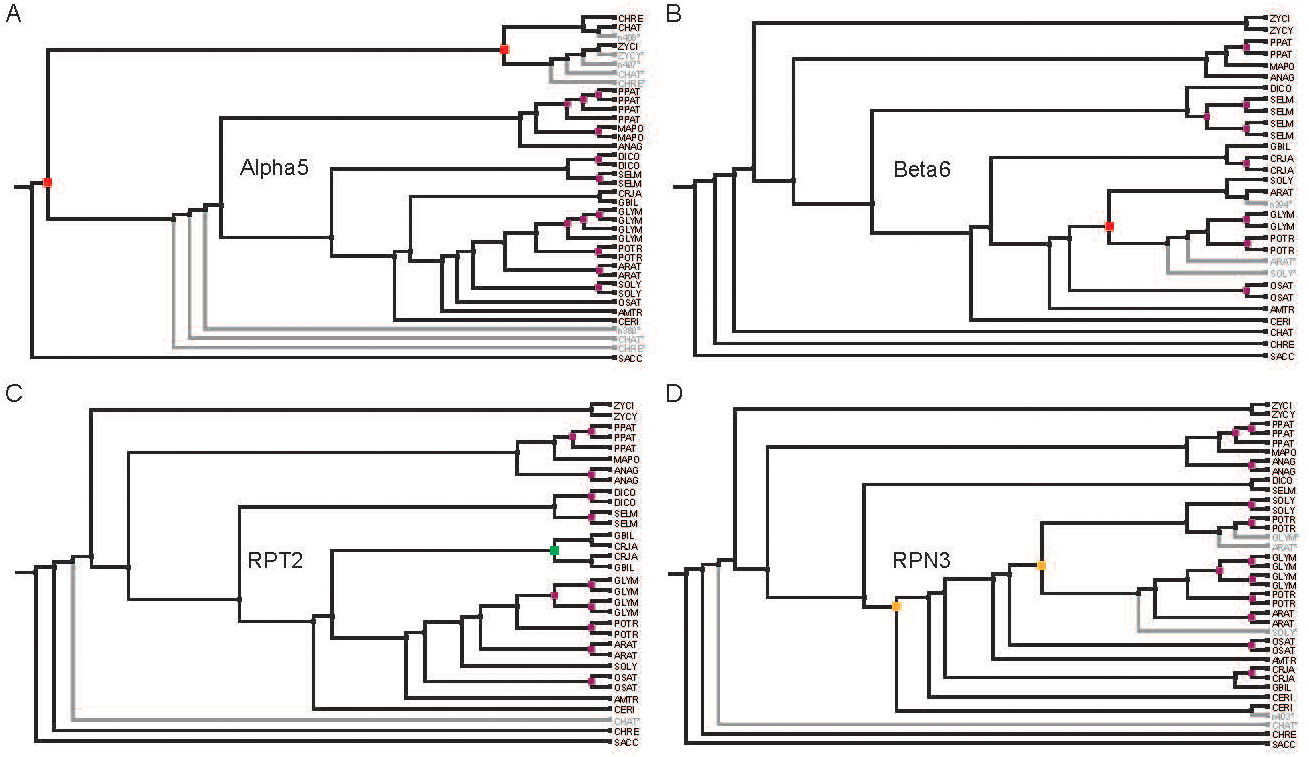
Reconciled trees illustrating different histories of proteasome subunit duplication. **A–D.** Black branches indicate extant genes. Gray branches indicate lost or absent genes. Red nodes indicate duplication events followed by loss of one duplicate in all descendant species. Orange nodes indicate duplication events followed by partial retention of duplicates. Green nodes indicate duplication events in which both duplicates were retained. Purple nodes indicate duplication events occurring after lineage divergence. **A.** Reconciled tree of Alpha5 subunit evolution. Two ancestral duplication events are predicted to have occurred early in *Viridiplantae* evolution, followed by loss of one duplicate in all descendant species. **B.** Reconciled tree of Beta6 subunit evolution. One angiosperm-specific duplication event is predicted to have occurred after the divergence of *O. sativa*, followed by loss of one duplicate in all descendant species. **C.** Reconciled tree of RPT2 subunit evolution. One gymnosperm-specific duplication event is predicted to have occurred, followed by retention of both duplicates. **D.** Reconciliated tree of the RPN3 subunit evolution, 1 event of Angiosperm-specific duplication has been predicted to happen after divergence of *O. sativa*, with conservation of the duplicates only in *P. tricocharpa*. 1 ancestral event of duplication has been predicted to happen around ferns divergence; which both duplicates are present in *C. richardii*.

**Figure S5.**
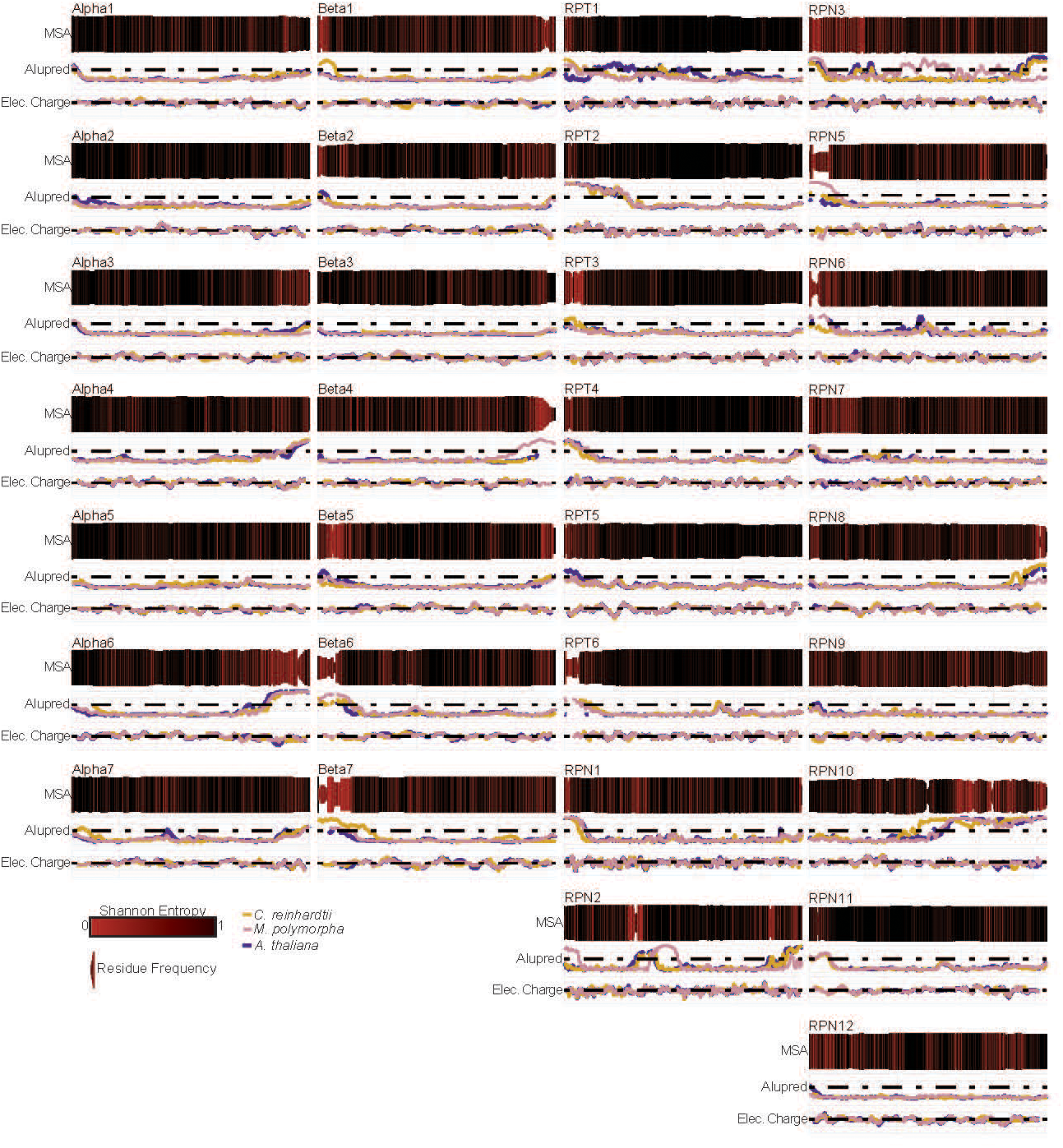
Summary of proteasome subunit sequence features from multiple-sequence alignments. For each 26S proteasome subunit family, codon-based multisequence alignments are represented. Alignment columns are colored by Shannon entropy; bar heights within columns indicate residue frequency at that position. First line plots show intrinsically disordered prediction for *C. reinhardtii*, *M. polymorpha* and *A. thaliana* representative subunits is represented; dotted line marks the IDR threshold. Second line plots show predicted local electrostatic charge for the representative subunits; dotted line marks neutral charge level.

**Figure S6.**
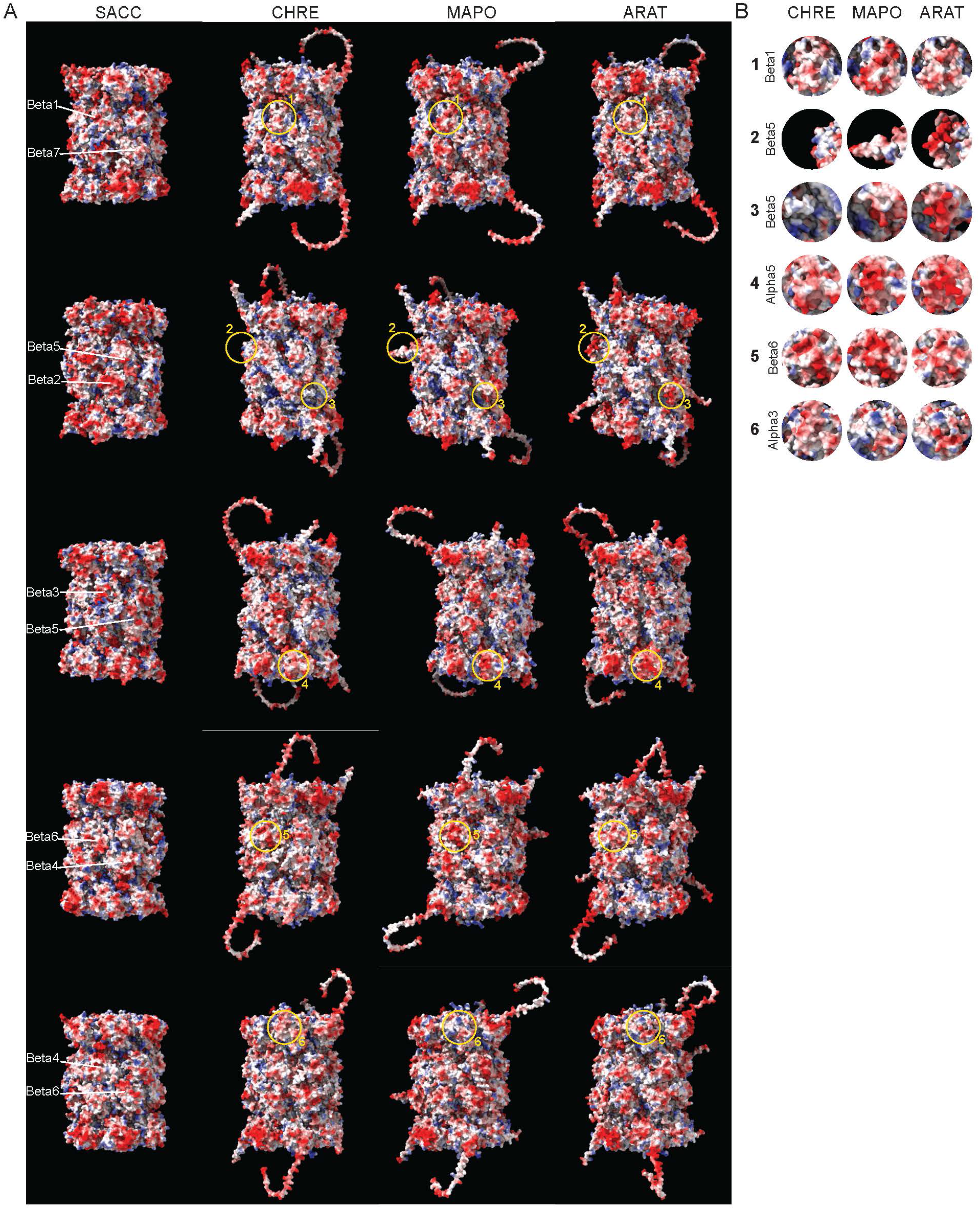
*In silico* reconstitution of 20S proteasome from *C. reinhardtii*, *M. polymorpha* and *A. thaliana*. **A.** Comparison of *in silico* reconstituted 20S proteasome complex from *C. reinhardtii*, *M. polymorpha* and *A. thaliana* with the *S. cerevisiae* crystal structure, shown from multiple views. On the S. cerevisiae complex, the β subunits facing the viewer are annotated. Surfaces are colored by electrostatic potential. Yellow circles indicate regions shown in panel B. **B.** Close-up views highlighting surfaces with notable differences in electrostatic potential.

**Figure S7.**
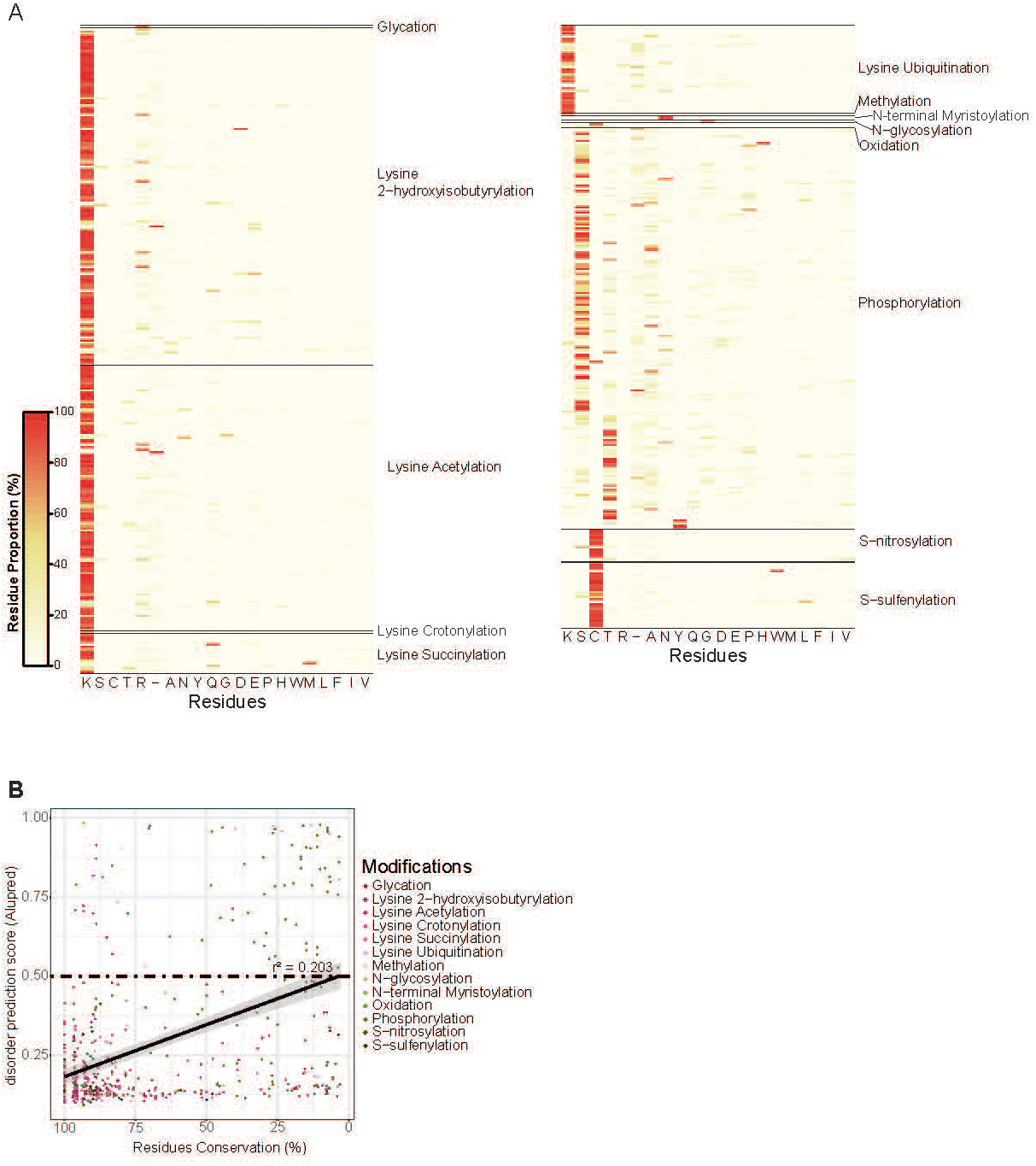
Polymorphism of post-translational modification sites in plant 26S proteasome subunits. **A.** Heatmap showing, for each alignment position bearing a reported PTM, the proportion of amino acid residues observed across species. **B.** Residue conservation versus average predicted IDR score (from Fig S4). Gray line indicates linear regression with R².

**Figure S8.**
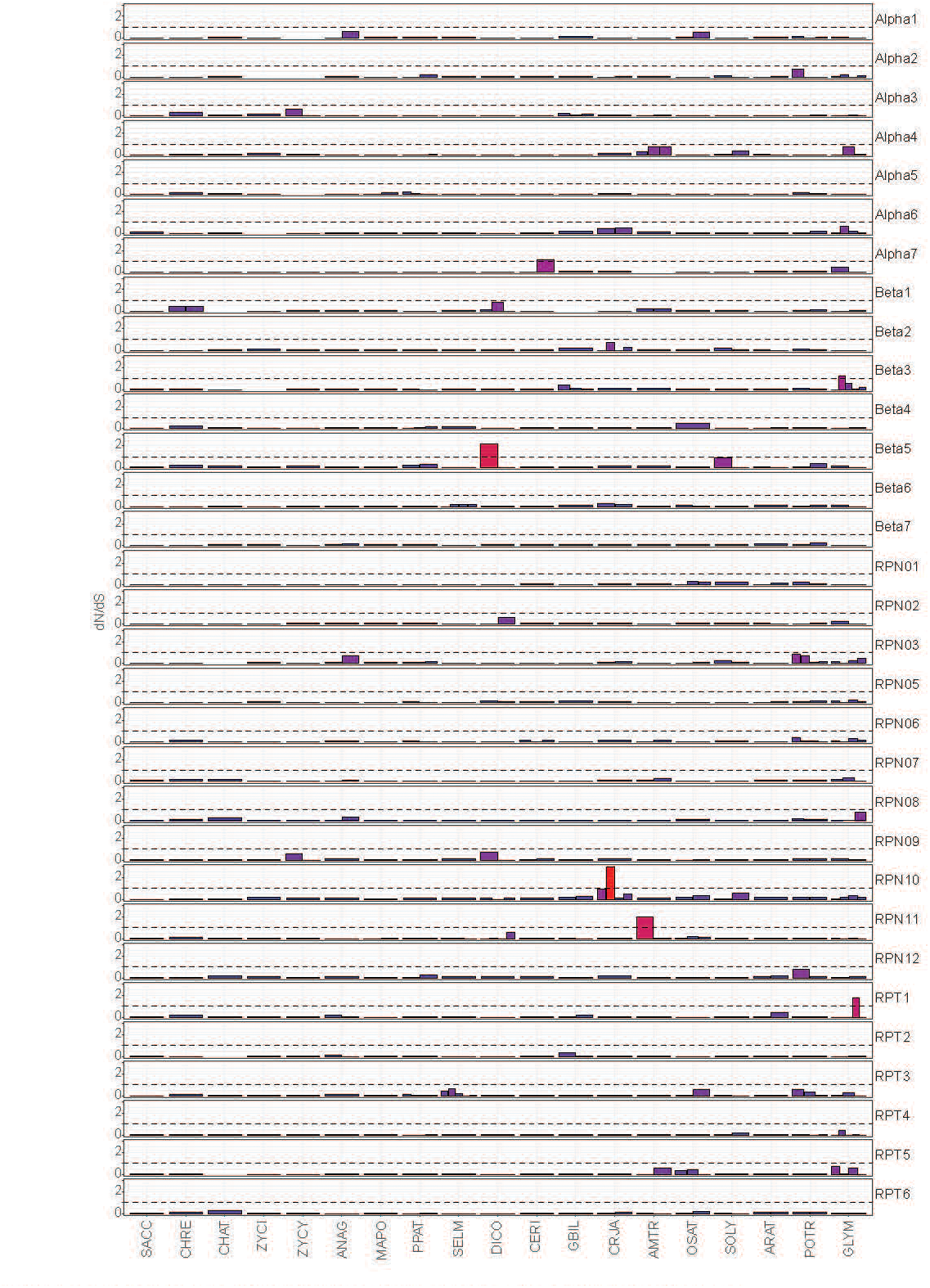
Positive selection analysis of 26S proteasome paralogs evolution. Bar plots summarize gene-specific dN/dS estimates for each proteasome gene on the terminal branch (branch after the last internal node). Bar colors encode the dN/dS value. The dotted line marks dN/dS = 1, above which positive selection is inferred.

**Figure S9.**
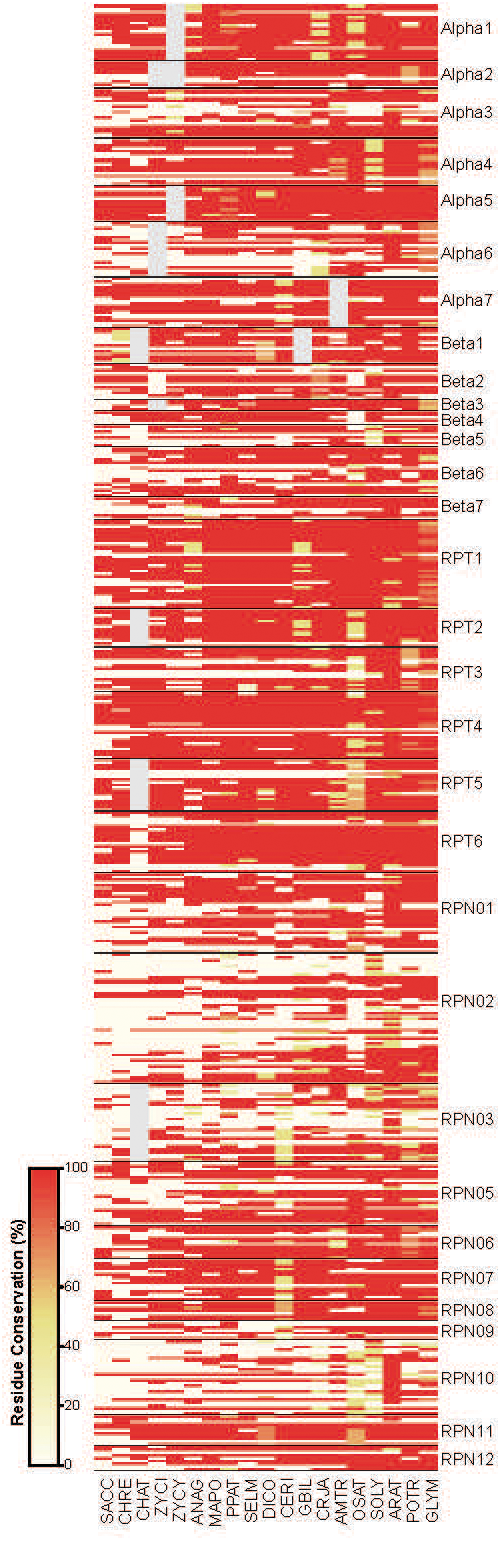
Conservation of post-translational modification site residues between paralogs. For each representative species and proteasome subunit family, residue conservation is shown for each identified post-translational modifications site. For species with paralogs, percent residue conservation at PTM sites is reported to indicate divergence or conservation between paralogs. Gray boxes indicate subunits absent from the genome.

**Figure S10.**
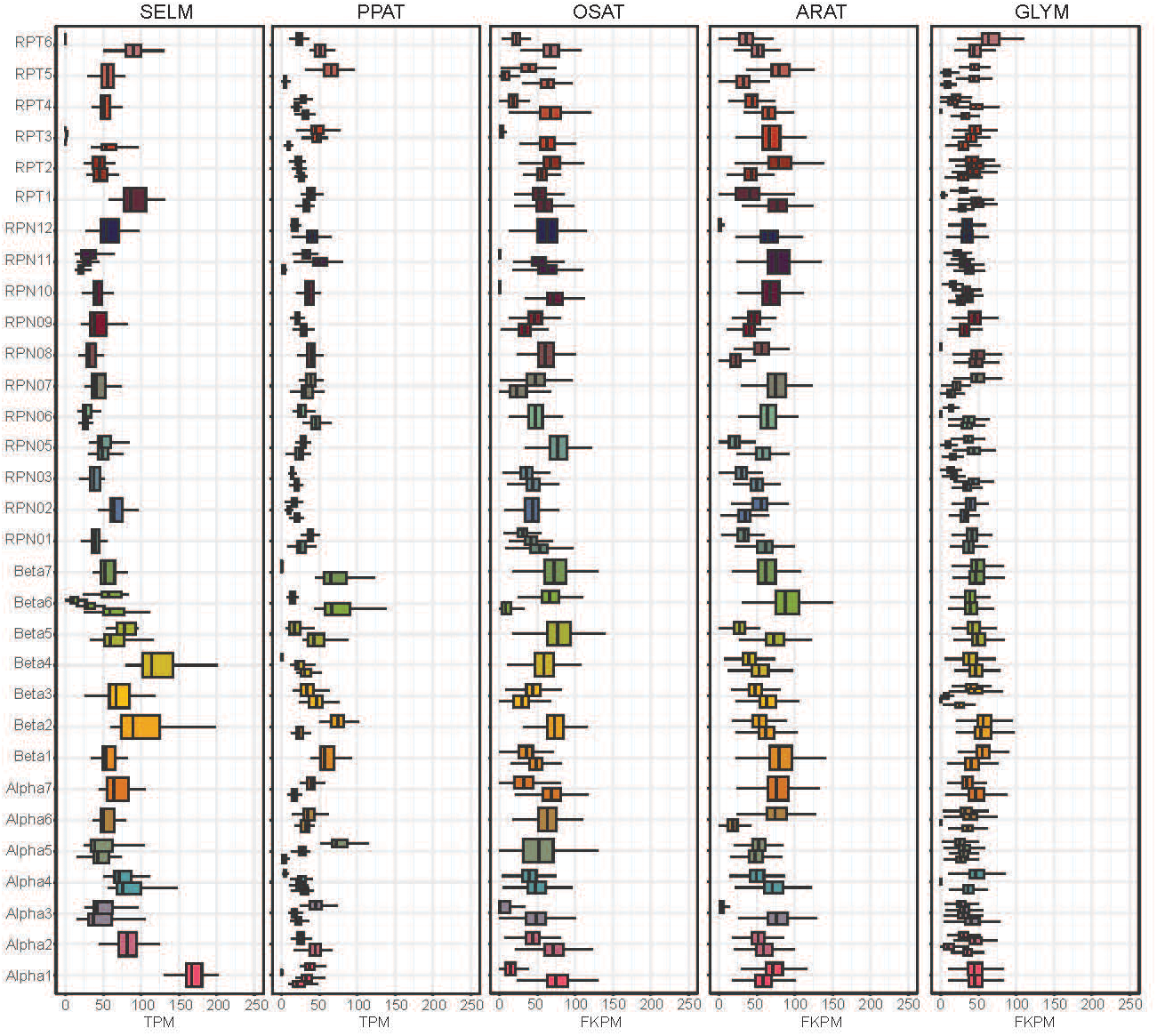
Proteasome paralogs mRNA abundance. Box plots of expression levels (TPM or FPKM) for proteasome genes in representative species with many duplications. Box plots are grouped and colored by subunit family.

**Figure S11.**
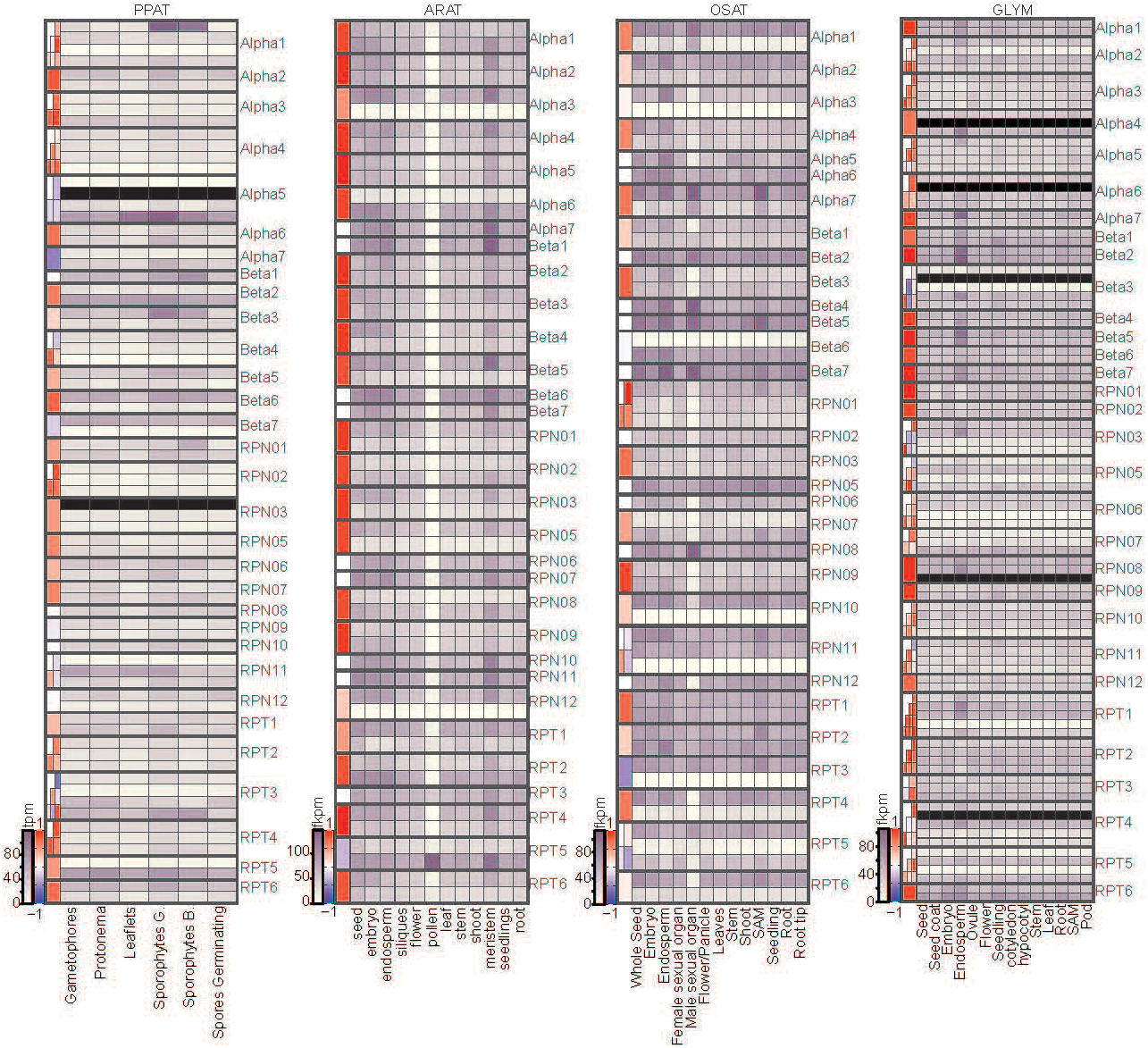
Analysis of expression level of 26S proteasome genes in developmental context. Heatmaps show expression levels (TPM or FPKM) for proteasome genes in representative species across developmental stages and/or tissues. Left-side panels display pairwise Spearman correlations between paralogs. Genes with no detectable expression are shown in black.

**Figure S12.**
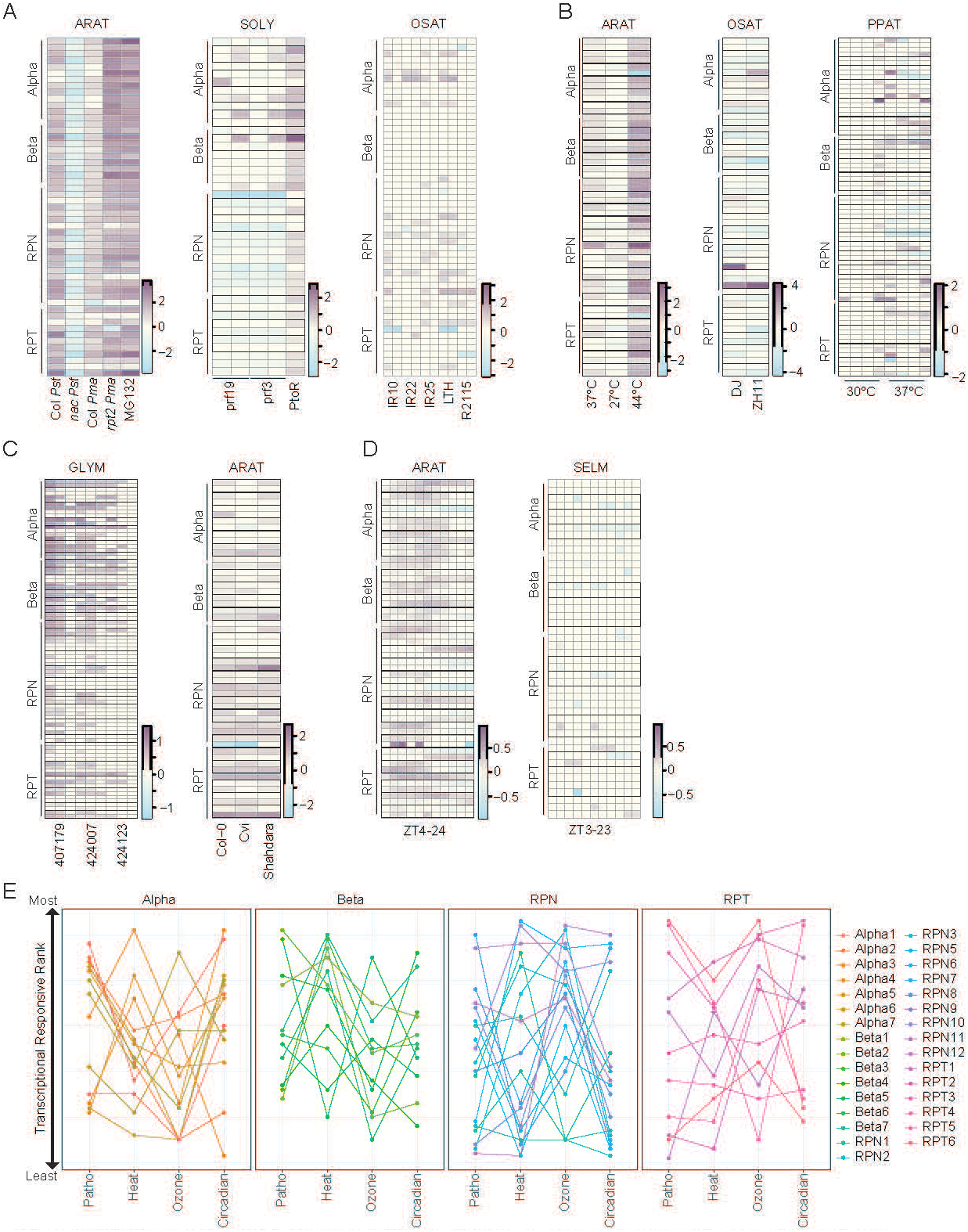
Expression fold-change of 26S proteasome genes across environmental contexts. **A.** Heatmaps of Log2 FC in expression during pathogen-related conditions in *A. thaliana*, *S. lycopersicum* and *O. sativa*. **B.** Heatmaps of Log2 FC under heat stress in *A. thaliana*, *O. sativa* and *P. patens*. **C.** Heatmaps of Log2 FC under ozone stress in *A. thaliana* and *G. max*. **D.** Heatmaps of Log2 FC across circadian times in *A. thaliana* and S*. moellendorfii*; the first time-point was used as the reference. **A-D.** Duplicated genes are outlined in black. The Log2 FC scale is shown at bottom right. Key environmental condition details are noted beneath each heatmap; see original studies for full descriptions. **E.** Dot plot representing the ranking of A. thaliana 26S proteasome genes from least to most responsive, based on the average absolute log2 fold change (log2FC) in each dataset shown in panels A–D. Dots are connected by lines to track their distribution across conditions. Dots and lines are colored according to subunit type; therefore, paralogs are shown in the same color.

**Figure S13.**
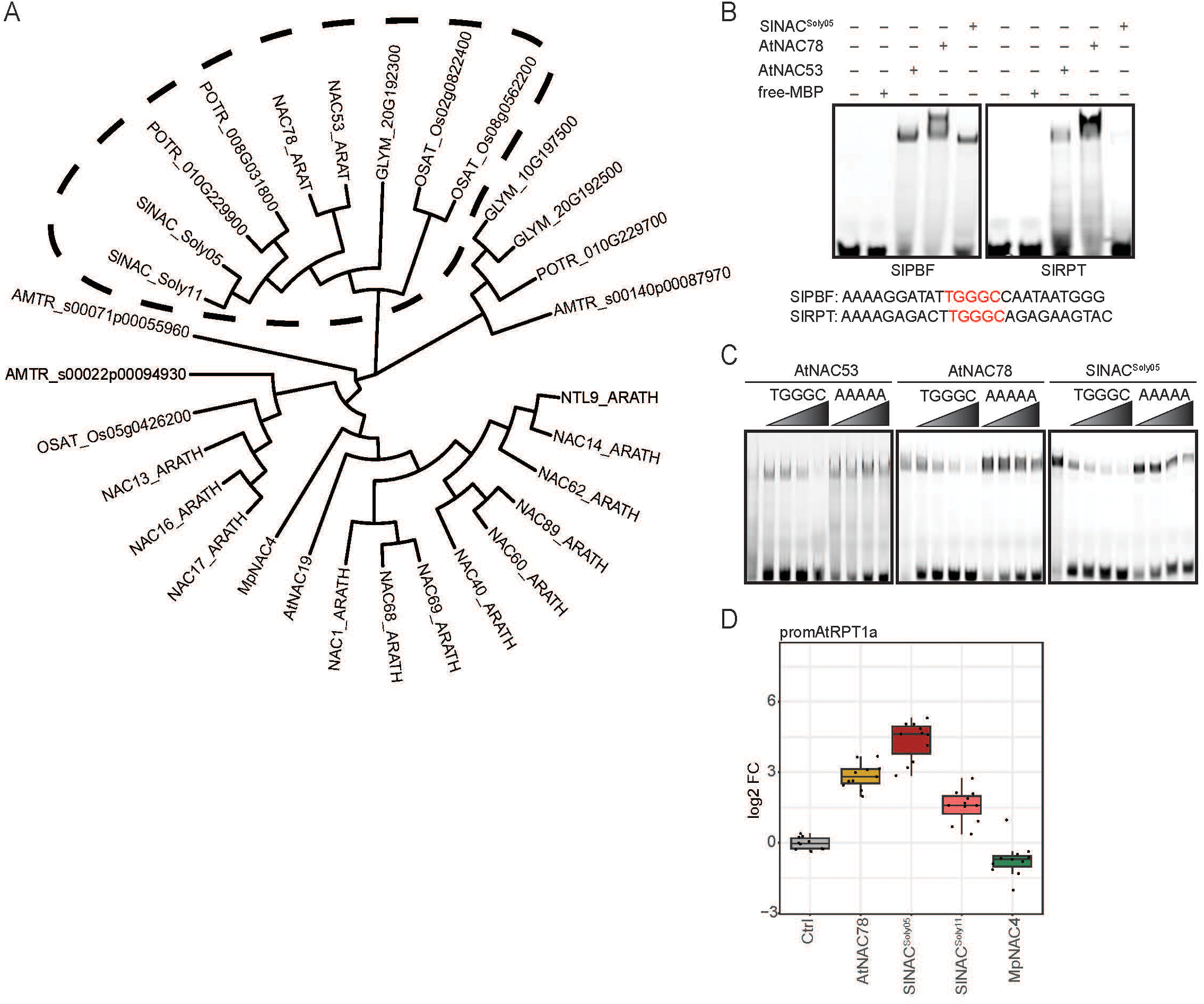
Conservation of NAC53/78-proteasome transcriptional regulation in *S. lycopersicum*. **A.** Hidden Markov model (HMM) phylogeny of NAC proteins related to AtNAC78; the NAC53/78 ortholog clade is outlined with a dotted line. **B.** EMSA with MBP-tagged AtNAC53/78, MBP-SlNAC^Soly05^, free-MBP using probes from *S. lycopersicum* proteasome genes promoters. Representative images from three independent experiments with similar results. **C.** EMSA competition using the SlPBF probe. Binding-induced shifts in the first lanes were challenged with increasing concentrations (10× to 100×) of unlabeled competitor probes: WT (TGGGC) or mutated (AAAAA). Representative of three independent experiments. **D.** Dual-luciferase promoter transactivation assay using the AtRPT1a promoter (-1 kb) in *A. thaliana* protoplasts expressing NAC78, SlNAC^Soly05^, SlNAC^Soly11^, MpNAC4 in *A. thaliana* protoplasts. Data are shown as Log2 fold change relative to control; pooled from three independent experiments.

**Figure S14.**
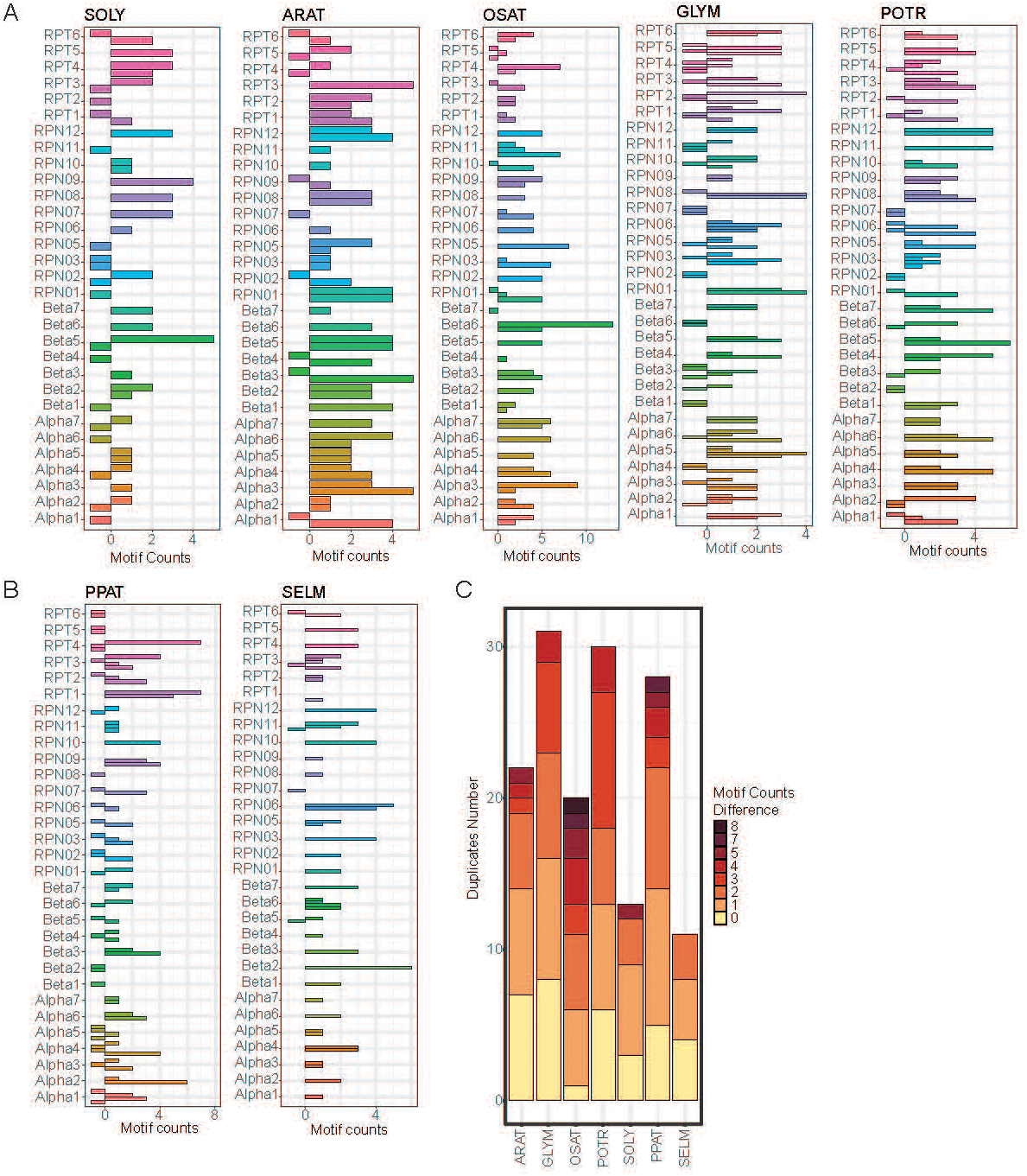
Analysis of proteasome-associated cis-elements enrichment in proteasome genes paralogs. **A.** Counts of [GA]GCCCA motifs in paralog promoters (-2 kb/+100 bp) from representative angiosperms. **B.** Counts of GC[TA]GC (*P. patens*) and AACCCTA (*S. moellendorffii*) motifs in paralog promoters (-2 kb/+100 bp). **A-B.** Bars are grouped and colored by subunit families; promoters with zero counts are shown as -1. **C.** Stacked bar plot of motif count differences between paralogs across all species analyzed in panels A-B.

**Figure S15.**
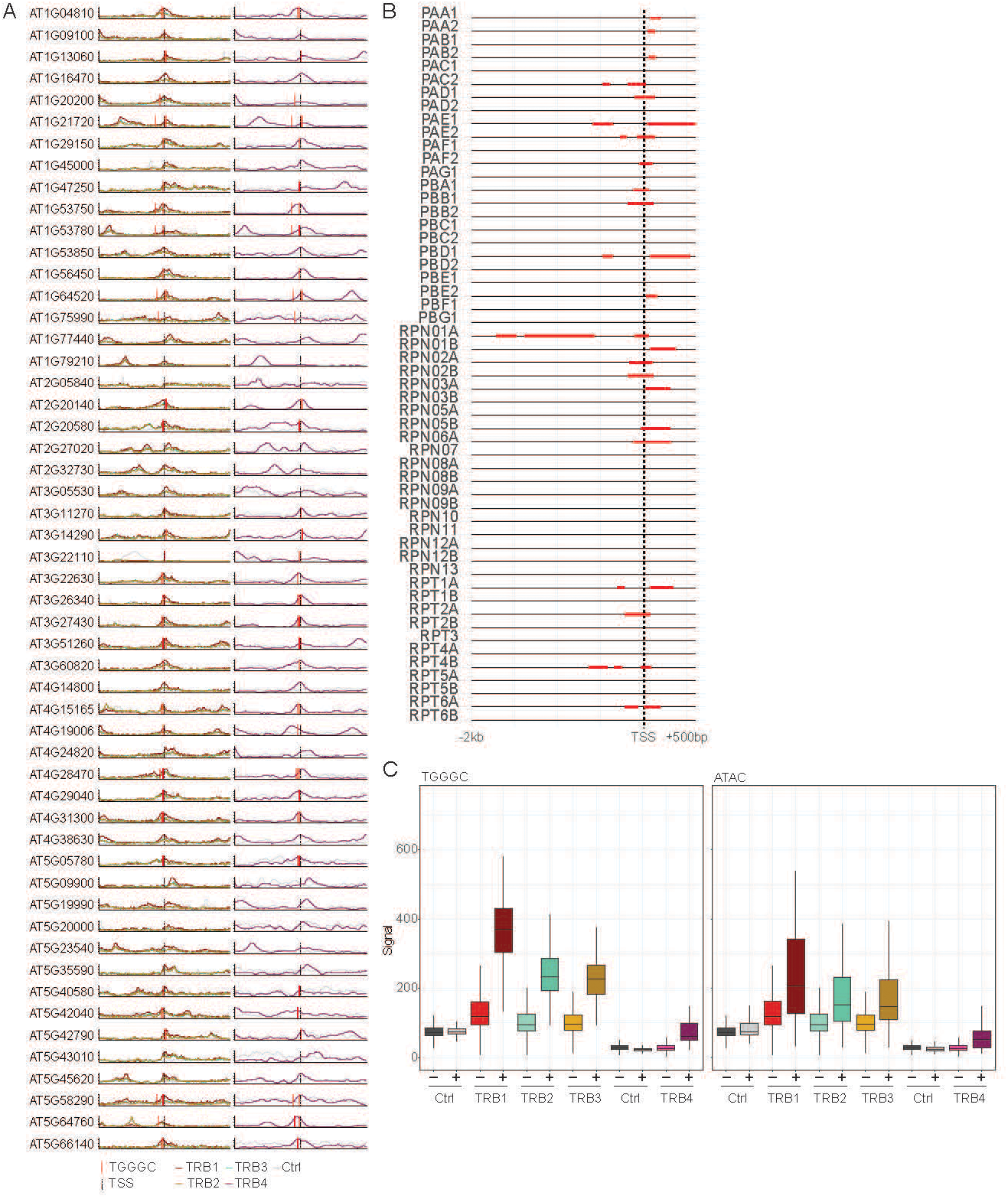
Analysis of AtTRBs association with 26S proteasome promoters. **A.** ChIP-seq signal for TRB1, TRB2, TRB3, and TRB4 across regions flanking 26S proteasome TSSs (-3 kb/+3 kb). Gray curves show controls; colored curves indicate individual TRBs. Dotted lines mark TSS positions; red ticks mark TGGGC motif sites. **B.** Proteasome promoters (-2 kb/+500 bp) with ATAC-seq-defined regions of significantly increased chromatin accessibility upon bacterial infection highlighted in red. Dotted line indicates the TSS. **C.** Comparison of TRB ChIP-seq signal over promoter segments bearing TGGGC elements or showing significant chromatin opening versus the remaining promoter regions of 26S proteasome genes.

**Figure S16.**
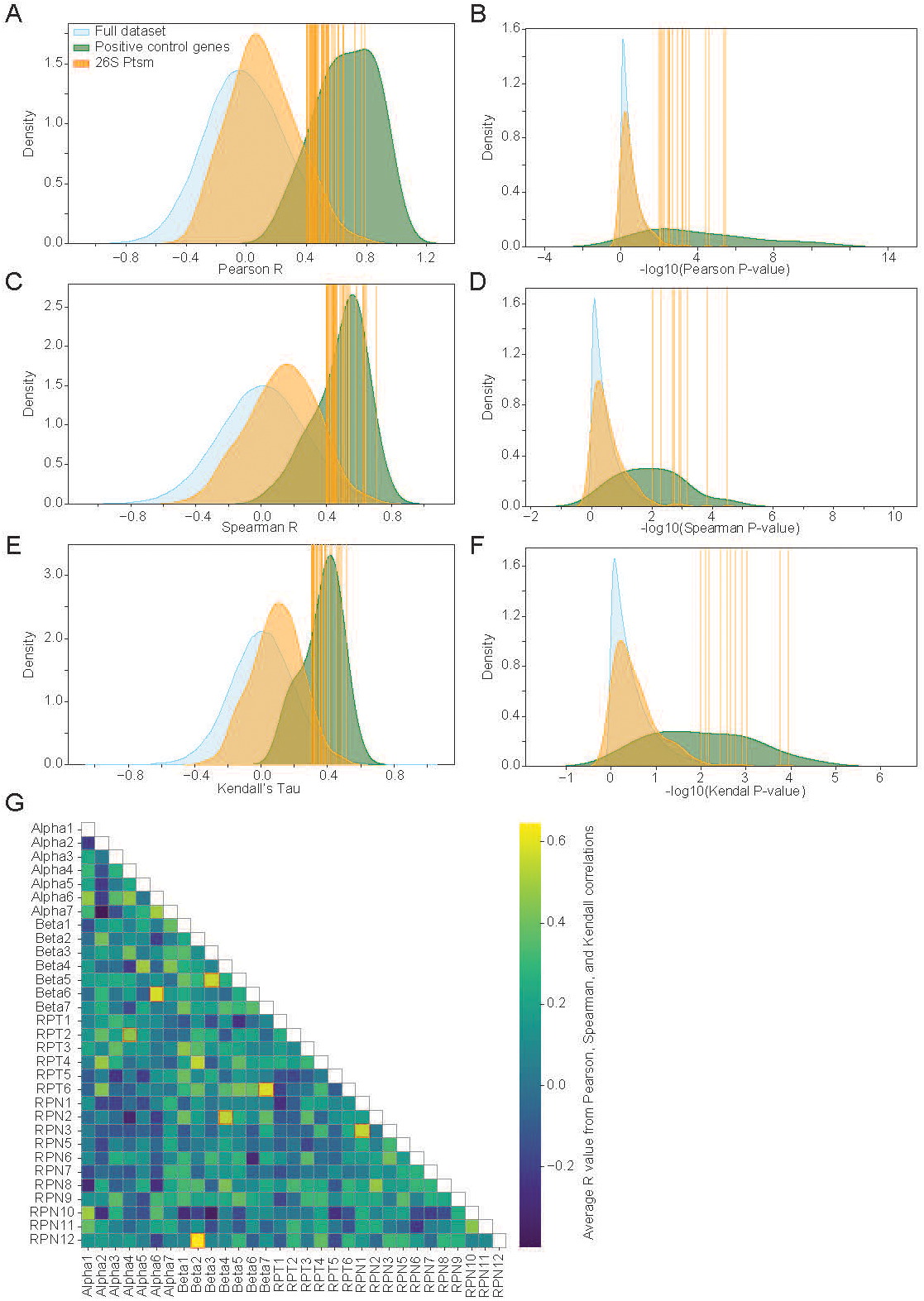
Evolutionary rate covariation among 26S proteasome subunits. **A-F.** KDE plots showing ERC R-values and negative log10 transformed P-values calculated with the Pearson (A, B), Spearman (C,D) and Kendall’s Tau (E,F) method. Blue distributions represent all pairwise gene combinations for the full dataset. Green distributions represent pairwise combinations of known subunits in the plastid caseinolytic protease (Clp) complex, chosen as a positive control. Orange distributions represent pairwise combinations of 26S proteasome subunits. Vertical lines indicate combinations for 26S proteasome subunits that exceed the statistical threshold: Pearson’s R > 0.4 (A), Pearson P-value < 0.01 (B), Spearman’s R > 0.4 (C), Spearman p-value < 0.01 (D), Kendall’s Tau > 0.3 (E), Kendall P-value < 0.01 (F). Correlations were calculated using Pearson’s correlation coefficient (A, B), Spearman’s rank correlation coefficient (C, D), and Kendall’s tau (E, F). Left panels show the distributions of correlation coefficients (A, C, E), whereas right panels show the corresponding significance values as −log10-transformed P-values (B, D, F). **G.** Pairwise evolutionary rate covariation between 26S proteasome subunits. The heatmap shows the average correlation coefficient obtained from Pearson, Spearman, and Kendall analyses for each pair of subunits. Red-outlined squares indicate subunit pairs that pass the statistical thresholds in all three correlation analyses.

**Figure S17.**
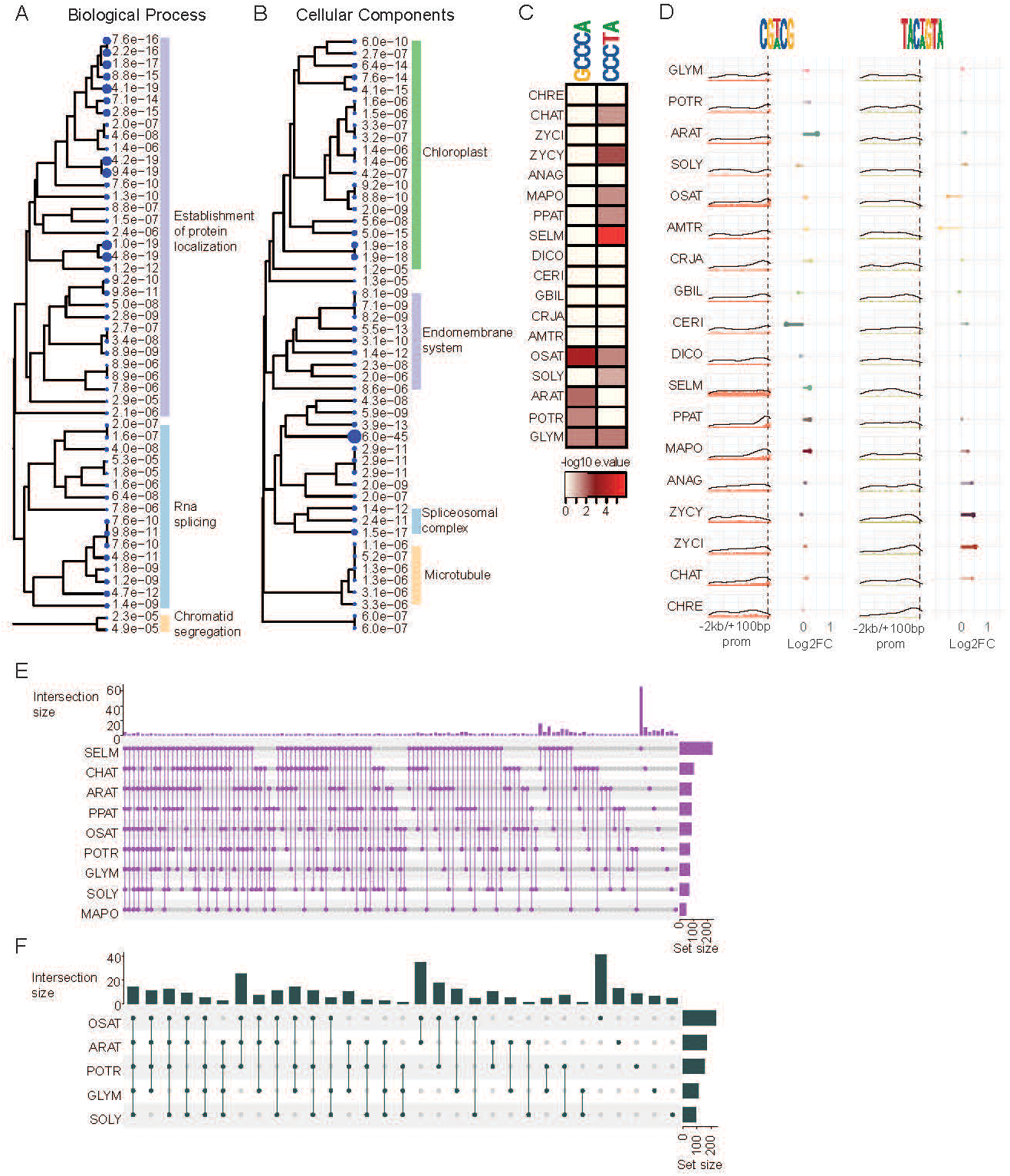
Proteasome-associated co-evolutionary protein network analyses. **A.** Cladogram of the top 50 enriched Gene Ontology Biological Process (GO:BP) terms among *A. thaliana* genes from the network in Fig. 5A. For clarity, specific terms are hidden and grouped into broader categories. **B.** Cladogram of the top 50 enriched Gene Ontology Cellular Component (GO:CC) terms among *A. thaliana* genes from the network in Fig. 5A. For clarity, specific terms are hidden and grouped into broader categories. **C.** STREME motif-enrichment summary for promoters of network genes across species. Based on motifs containing GCCCA or CCCTA, -log10(e-value) is shown per species. **D.** Enrichment and positional analyses of CGWCG and TACWGTA motifs in promoters (-2 kb/+100 bp) of network genes across species. Bar plots show motif positional frequency; dotted lines mark the TSS. Black curves represent density functions. Lollipop charts display genome-wide enrichment as log2 fold change; lollipop size indicates - log10(p). Fisher’s exact test was used; lollipops are semi-transparent where p > 0.001. **E.** Intersection plot of network genes bearing AACCCTA in their -400 bp/+100 bp promoter window for the nine species in which this motif is significantly enriched. **F.** Intersection plot of network genes bearing [GA]GCCCA in their -400 bp/+100 bp promoter window for the five species with significant enrichment.

**Table S1. Information on public datasets used in this study**

**Table S2. Plant 26S proteasome gene identifiers and sequences**

**Table S3. Top 10 candidate motifs and associated scores for 26S proteasome promoters in each species analyzed**

**Table S4. Post-translational modification sites identified and analyzed in 26S proteasome subunits across plant species**

**Table S5. Significant evolutionary rate covariation with proteasome subunits, including functional annotations of network members**

